# Programmed chromosome elimination correlates with the overexpression of cohesin and additional B chromosome-encoded genes in *Aegilops speltoides*

**DOI:** 10.64898/2026.07.20.739560

**Authors:** Gihwan Kim, Jianyong Chen, Johannes Thiel, Amanda Souza Camara, Daiyan Li, Tereza Bojdová, Jörg Fuchs, Maria Cuacos, Veit Schubert, Mohammad-Reza Hajirezaei, Srijan Jhingan, Axel Himmelbach, Bhanu Prakash Potlapalli, Jochen Kumlehn, Jędrzej Szymański, Anne Fiebig, Miroslava Karafiátová, Jan Bartoš, Andreas Houben

## Abstract

Programmed chromosome elimination is a highly controlled developmental process in which specific chromosomes are selectively lost from defined cell types during development. Despite its broad occurrence across plants and animals, the molecular mechanisms driving the tissue-specific elimination of chromosomes remain largely unresolved. Here, we exploit the root-specific elimination of supernumerary B chromosomes in *Aegilops speltoides* as a tractable model to identify the genetic basis of programmed chromosome loss. A high-quality, chromosome-scale genome assembly was generated, assigning 398 Mb of sequence to the *Ae. speltoides* B chromosome. Transcriptome profiling across seven tissue types representing chromosome elimination-active, elimination-negative, and B chromosome nondisjunction conditions identified 3,262 genes consistently upregulated in elimination-associated tissues, including 1,035 B genes. Stepwise subtraction of genes expressed in post-elimination and B chromosome-retaining reference tissues, followed by intersection with genes expressed during B nondisjunction in anthers, identified a candidate gene set enriched for chromosome segregation functions. From this set, we prioritized *SYN2-B*, a B chromosome-encoded cohesin α-kleisin subunit whose *Arabidopsis thaliana* ortholog *AtSYN2* induces chromosome bridges and micronucleus formation when misregulated. *CENH3-B*, an α-type centromeric histone variant identified through GO enrichment analysis of B genes expressed in elimination-associated tissues, was shown to be incorporated in centromeres of both A and B chromosomes by transient gene expression assays using protoplasts and 3D structured illumination microscopy. These findings support a model in which B chromosome-encoded perturbations of cohesin activity and centromere composition contribute to selective B chromosome nondisjunction and their elimination in root tissues. Moreover, the SYN2-B promoter is enriched for ethylene response factor-binding sites compared to its A-encoded paralog, suggesting that ethylene is implicated in the root identity pathway driving root-specific B chromosome elimination.

## Introduction

In most organisms, the genetic information remains constant in all cells throughout the life cycle. However, there are exceptions where the elimination of specific DNA is part of the developmental program. Elimination can range from small DNA fragments to entire chromosomes and has been documented across eukaryotes, from unicellular ciliates to animals and plants ^1, 2^. This process takes place in somatic cells, while the germline genome remains intact in most species. A variety of hypotheses have been proposed to explain the significance of programmed DNA elimination, including mechanisms of sex determination, gene dosage compensation, gene silencing, germline development, and germline and soma differentiation ^3^. Considering its wide phylogenetic distribution, programmed DNA elimination probably evolved independently in different lineages. A key question in DNA elimination is how chromosomes and sequences are selectively lost ^4^.

Typical examples of chromosome undergoing elimination are germline-restricted chromosomes (GRC) described in hagfish and lampreys ^5, 6, 7, 8^, passerine birds ^9, 10, 11^ and dipteran families ^12, 13, 14, 15, 16, 17^. In addition, B chromosomes (Bs) can be eliminated from specific tissues in certain species.

B chromosomes are dispensable chromosomes that occur in addition to the standard A chromosomes (As) ^18, 19^. Although not required for normal development, Bs can persist by exploiting host chromosome segregation machinery and increasing their own transmission ^19, 20, 21^. However, tissue specific elimination of Bs has been reported in several plant species, including *Aegilops mutica, Aegilops speltoides, Sorghum purpureosericeum, Xanthisma texanum, Agropyron cristatum* and *Poa alpina* ^22, 23, 24, 25, 26^ as well as in animal species, including *Nasonia vitripennis*, *Astyanax mexicanus*, *Locusta migratoria* ^27, 28, 29, 30^. The process of B chromosome elimination is likely more common than documented, as chromosome number and genome size studies frequently rely on analysis of a single tissue type.

The goatgrass *Aegilops speltoides* provides a striking example of tissue-specific B chromosome elimination. Its Bs are stably maintained in above-ground tissues but eliminated in roots ^22, 31^. Individual plants can carry up to eight identical Bs ^22^. In contrast, in wild sorghum (*Sorghum purpureosericeum*), B chromosome elimination is nearly complete in most mature somatic tissues ^32, 33^. In *Ae. speltoides*, B chromosome elimination begins at the onset of embryonic differentiation in the proto-root region and proceeds through mitotic B chromatid nondisjunction, anaphase lagging, micronucleation, and degradation ^31^. Notably, despite this unusual lagging of Bs, cell cycle progression remains largely unaffected. Whether the mitotic spindle assembly checkpoint is impaired or whether Bs escape the checkpoint control remains unknown.

Lagging Bs of *Ae. speltoides* retain centromere-associated features, including the presence of the centromere-specific histone H3 variant (CENH3/CENPA) and microtubule attachment, indicating that complete loss of centromere identity is not responsible for their elimination ^31^. The elimination process is chromosome-type specific, because in wheat x *Ae. speltoides* F1 hybrid plants carrying both rye and *Ae. speltoides* Bs, only the *Ae. speltoides* B chromosome is eliminated from roots, whereas the rye B chromosome is retained ^31^. The absence of B-A translocation chromosomes possessing a derived B centromere in root cells suggests that the B chromosome centromere itself is a key component of the chromosome elimination process ^34^.

Comparing chromosome elimination and post-meiotic chromosome drive in *Ae. speltoides* reveals striking differences and similarities in their cellular mechanisms ^31, 35^. In both processes, A chromatids separate during anaphase, whereas B chromatids undergo nondisjunction, likely due to a delayed release of B sister chromatid cohesion. However, the spindle symmetry differs between the two processes. During the first pollen mitosis, cell division is asymmetric, whereas the spindle in roots is symmetrical. As a consequence, lagging Bs form micronuclei and undergo elimination in root cells. In contrast, the asymmetric spindle geometry at the first pollen mitosis, allows the inclusion of the lagging joined B chromatids into the generative nucleus, resulting in B chromosome drive. Our pilot transcription analysis, comparing plants carrying B chromosomes (+B) with those lacking them (0B) across a single tissue type, identified ∼2,900 up-regulated unigenes associated with B chromosome elimination ^36^. Gene ontology enrichment analysis revealed overrepresentation of chromosome-, centromeric region-, and chromosomal region-associated terms in the cellular component category. However, the molecular basis of root-specific B elimination has remained unresolved. Understanding the molecular mechanisms underlying the dysfunction of tissue- and chromosome-type-specific cohesion may provide insights into the process of chromosome nondisjunction, a major cause of genetic disease across species ^37, 38^.

In this study, we assembled the sequence of the *Ae. speltoides* B chromosome. To identify B chromosome-encoded candidate genes potentially involved in chromosome elimination, we performed an extended, genome-resolved transcriptomic analysis. This approach revealed that B-encoded genes are significantly enriched for functions related to chromosome segregation, highlighted by a B-specific cohesin subunit (*SYN2*) and a *CENH3* variant. Our functional analyses demonstrate a role for *SYN2* activity in somatic nondisjunction, suggesting that *SYN2-B* acts as a *trans*-acting candidate factor associated with B chromosome elimination. Furthermore, the *SYN2-B* promoter is enriched with ethylene response factor-binding sites compared to its A-encoded paralog. This finding suggests that an ethylene-influenced, root-identity pathway may activate the root-specific program of B chromosome elimination.

## Results

### A hydroponic induction system for *Ae. speltoides* provides isogenic B-containing and B-depleted tissues for sequence analysis

The annual outcrossing species *Ae. speltoides* shows high genetic diversity ^39^, and its A and B chromosomes are highly similar in sequence ^31^. To limit confounding variation and improve the identification of B-associated sequences, we restricted our analysis to a single plant. To obtain sufficient material for the analysis, we established an induction system for adventitious roots (ARs) that enabled the hydroponic clonal propagation of a single *Ae. speltoides* plant harboring three B chromosomes in aerial tissue (Supplementary Fig. 1). ARs are specialized roots that emerge from non-root parts of a plant, e.g. stems ^40^. Application of an iron-supplemented solution (0.01 mM Fe-EDDHA, 1.4 mM CaSO₄, 1 mM MES, pH 5.8) as described in Hilo, Shahinnia ^41^ stimulated the growth of ARs at stem cuttings and allowed the vegetative propagation of plants (Supplementary Fig. 2).

Our previous study revealed partial B presence in scattered cell lineages of young Ars ^31^. To determine when the B elimination process in ARs is complete, flow cytometric analysis was performed. AR buds approximately 1 mm in size showed, in addition to the main nuclear peaks representing 2C and 4C nuclei without B chromosomes, additional nuclear populations with increased DNA content, consistent with the presence of B-containing nuclei (Fig. 1a). In contrast, AR-derived lateral roots displayed only 0B-equivalent 2C and 4C peaks, indicating that B elimination was complete in this tissue (Fig. 1a, b, Supplementary Fig. 3). To determine the spatial progression of B elimination in developing ARs, 50 µm tissue sections of 1 mm AR buds were analyzed by FISH using A and B chromosome-specific repeats as probes (Fig. 1c, d). B-specific signals were largely absent from the AR tip, whereas they remained detectable in the root-stem transition zone (Fig. 1d). The epidermis covering developing AR buds also retained B-specific signals, consistent with its origin from B-carrying stem tissue (Fig. 1d). B-positive micronuclei were also observed in this region, indicative of ongoing B chromosome loss (Fig. 1e). Together, these data show that B elimination progresses during AR development and that AR-derived lateral roots provide B-depleted tissue from the same clonal +B plant.

**Fig. 1.**
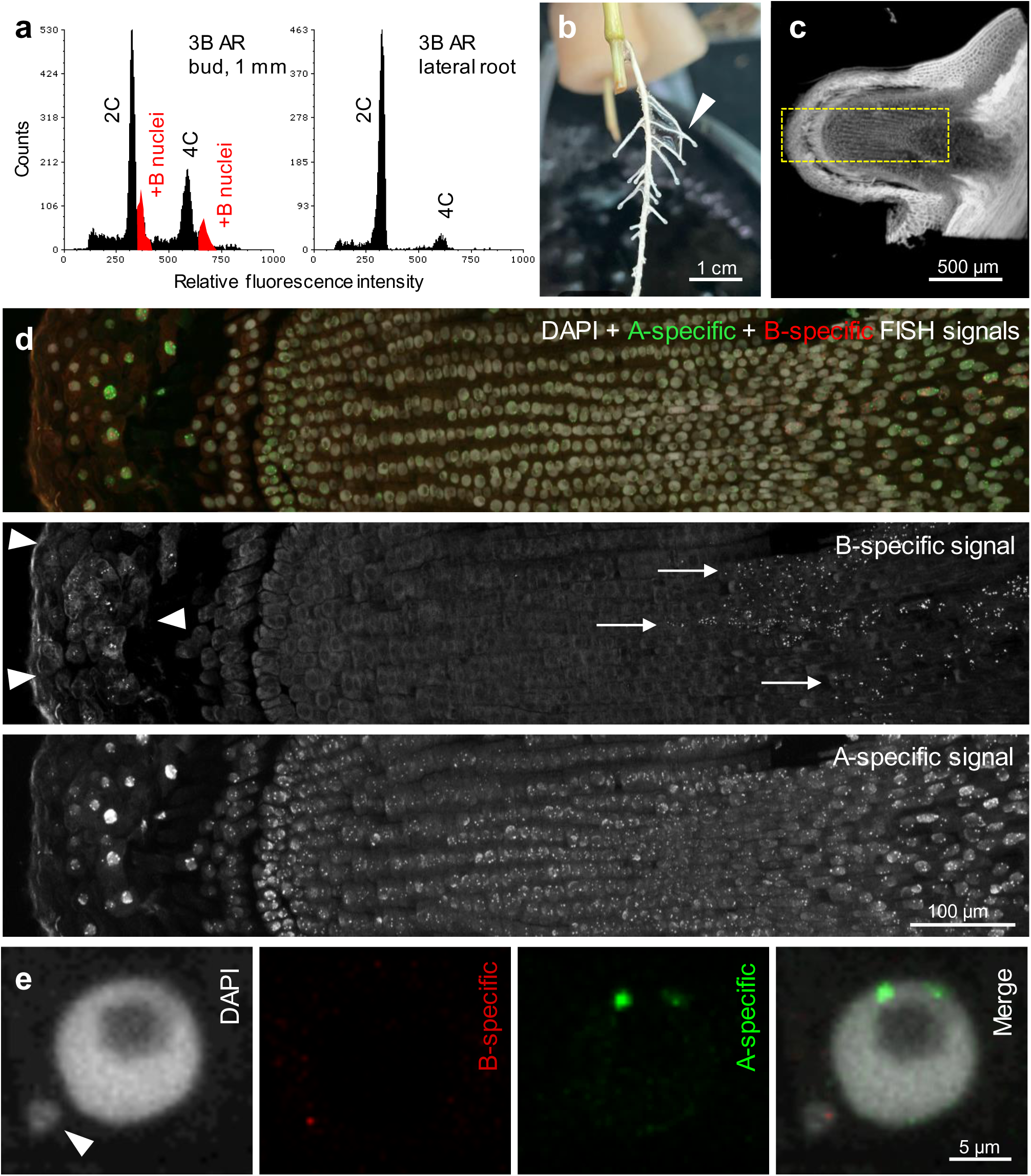
B chromosome elimination during adventitious root development in *Ae. speltoides*. (**a**) Flow-cytometric DNA content profiles of a 1 mm adventitious root (AR) bud and AR-derived lateral root from a plant carrying 3B. AR bud showed 0B-equivalent 2C and 4C peaks together with additional B-containing nuclear populations, whereas the AR-derived lateral root showed only 0B-equivalent 2C and 4C peaks. (**b**) Representative AR-derived lateral roots (arrowheaded) used as B-free root tissue. (**c**) Representative 50 µm tissue section of a young AR bud used for FISH. (**d**) FISH analysis of the tissue section shown in (c) using the A chromosome-specific repeat pBs301 and the B chromosome-specific repeat AesTR183. Arrows indicate B-specific signals in the root-stem transition zone. B-specific signals were absent from the AR tip, whereas B-specific signals remained detectable in the root-stem transition zone, indicating spatial progression of B chromosome elimination. Arrowheads indicate B chromosome-positive epidermis. (**e**) Magnified view of a cell with a B-positive micronucleus (arrowed) after FISH detected in the root-stem transition zone.

### Whole-genome assembly of *Ae. speltoides* with B chromosomes

A first sequence of the *Ae. speltoides* B chromosome has been available since 2020 ^31^, but this assembly represents ∼16% of the B only and thus cannot be used to uncover B-encoded genes. Now, high-molecular-weight DNA from +B leaf tissue of the clonally propagated +B plant was used to generate a chromosome-scale genome assembly. A total of 154 Gb of PacBio HiFi reads and 36.4 Gb of Nanopore reads (>25 kb) were generated for primary assembly using hifiasm ^42^. The resulting 5.47 Gb assembly achieved 93.7% BUSCO completeness (contig N50 = 17.1 Mb, Table 1).

**Table 1.** Statistics of the *Ae. speltoides* +B genome assembly by QUAST and BUSCO.

|  |  |  |
| --- | --- | --- |
| **QUAST** | Number of contigs | 1794 |
|  | Largest contig | 95 Mb |
|  | Total length | 5.47 Gb |
|  | N50 | 17.1 Mb |
|  | N75 | 8.0 Mb |
|  | Mismatches # N's | 0 |
| **BUSCO** | Total BUSCO groups searched | 3236 |
|  | Complete BUSCOs | 3032 (93.7%) |
|  | Missing BUSCOs | 42 (1.3%) |

Approximately 104 Gb of Hi-C sequencing data derived from leaf tissue of the same +B plant were employed to scaffold the primary contigs. The assembly yielded eight large scaffolds (398–835 Mb), each displaying a characteristic Rabl configuration (Supplementary Fig. 4a). Alignment of these scaffolds to the reference genome of *Ae. speltoides* accession AEG-9674-1 without B chromosome ^43^ revealed that seven of the eight scaffolds showed strong synteny with the standard A chromosomes 1S–7S (Supplementary Fig. 4b). To determine whether the remaining large scaffold corresponded to the B chromosome, we generated ∼60 Gb of whole-genome sequencing (WGS) data from 0B AR-derived lateral root tissue of the same plant, as well as approximately 36 Gb of WGS data from +B leaf tissue. Comparative read-mapping analyses showed that the eighth scaffold exhibited normal sequencing coverage in +B leaf-derived data but substantially reduced coverage in 0B AR-derived data (Supplementary Fig. 4c). Thus, the 398 Mb scaffold represents the B chromosome, accounting for 69% of its size as estimated by flow cytometry (Supplementary Fig. 4c). Additionally, 4.81 Gb of contigs were assigned to the seven A chromosomes, representing 91% of their estimated size (1C=5.27 Gb). Consequently, we produced a high-quality chromosome-scale assembly of *Ae. speltoides* carrying B chromosomes.

### Genome-guided transcriptome profiling reveals B-encoded genes associated with chromosome segregation

To identify genes associated with the B chromosome elimination process, RNA-seq was performed across various developmental stages and different tissues in which B chromosome behavior differs. RNA was isolated from whole embryos at 6 to 8 days after pollination (DAP) and 22 to 25 DAP, from laser-capture microdissected (LCM) embryonic roots (Supplementary Fig. 5), from AR buds (1 mm), primary roots (3 days after germination (DAG)), shoots (3 DAG), and first pollen mitosis (PM I) anthers from 0B and +B plants (Supplementary Table 1). This sampling design captured tissues associated with root-specific B elimination, a B-retaining reference tissue (shoots), a complete B-loss reference tissue (primary roots), and a tissue undergoing B-nondisjunction-associated drive (PM I anthers) for comparative analysis. Using all +B RNA-seq datasets, we annotated the *Ae. speltoides* genome assembly containing the B chromosome. This annotation identified 59,981 transcripts and 47,792 protein-coding genes on the seven A chromosomes and 5,940 transcripts and 4,196 protein-coding genes on the B chromosome (Supplementary Table 2-3).

Principal component analysis revealed tight clustering of biological replicates, with PC1 (34.4%) separating samples primarily by tissue identity and B chromosome presence, with +B and 0B samples showing clear separation across most tissues. PC2 (21.1%) captured additional transcriptomic variation, particularly resolving subtle differences within tissue groups (Supplementary Fig. 6). The embryos, embryonic roots, as well as AR buds were included as B elimination-associated tissues at early developmental stages, whereas primary roots represented later root development after B elimination. Shoots were included as a B-retaining control tissue, and PM I anthers were included as a tissue associated with B nondisjunction. PCA confirmed the quality of the RNA-seq dataset and indicates that the presence of B chromosomes significantly affects the transcriptomes of different tissues.

Differential expression analysis between +B and 0B samples identified genes consistently upregulated in tissues undergoing B elimination. Genes upregulated in +B across all four B elimination-associated tissues were intersected to define a core set of 3,262 genes, of which 1,035 were B-encoded genes (Fig. 2a). Subtraction of genes also upregulated in primary roots of the +B plant, where Bs are already absent, reduced this set to 1,850 genes (including 949 B genes).

**Fig. 2.**
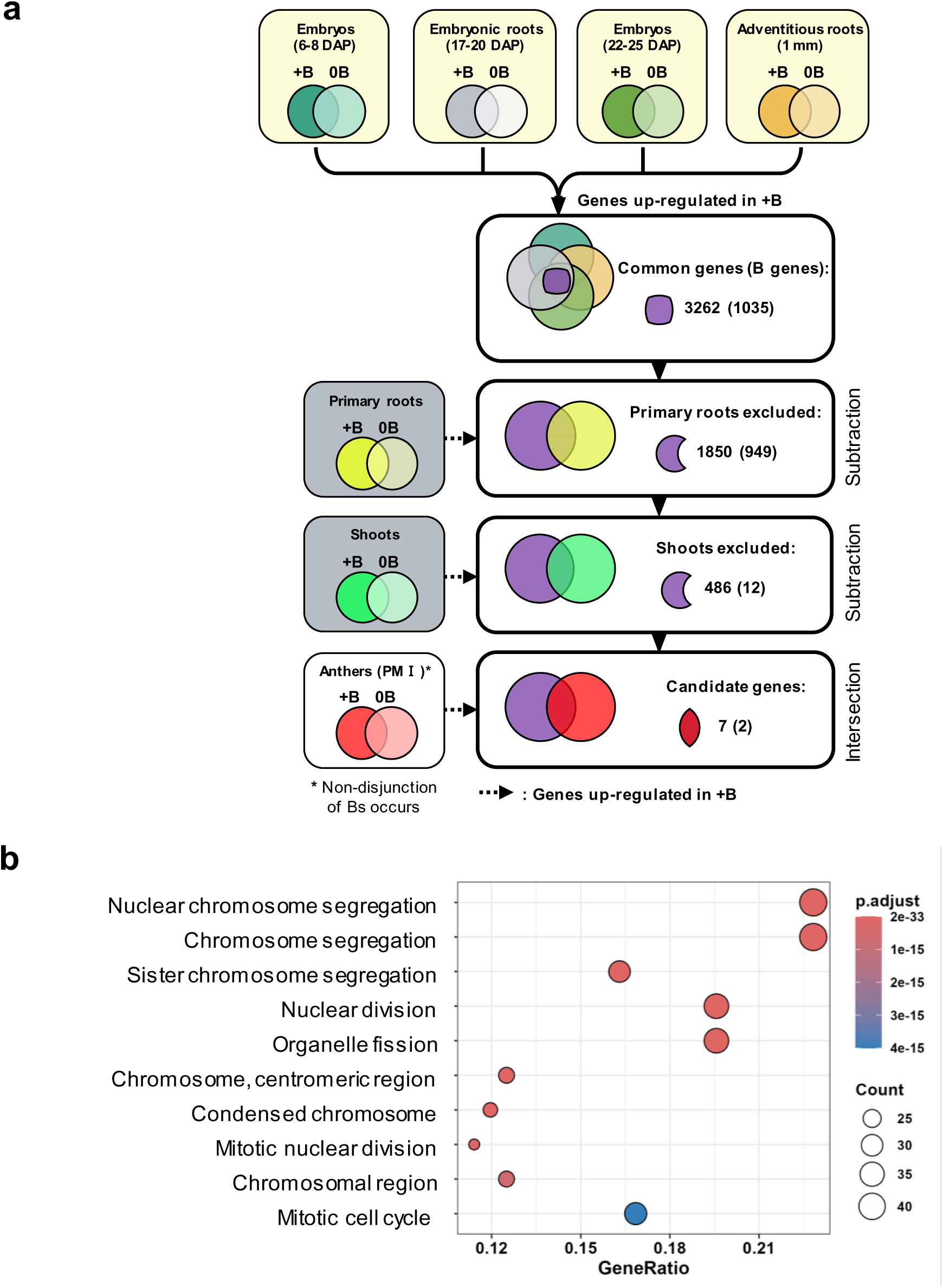
Comparative RNA-seq strategy and stepwise candidate gene prioritization. (**a**) Comparative RNA-seq strategy to identify genes associated with the B chromosome elimination process. Tissues undergoing B elimination are shown in yellow boxes. Tissues in which B elimination is not observed are shown in gray boxes. Venn diagrams illustrate genes upregulated in +B relative to 0B samples across tissues. Stepwise subtraction of primary root- and shoot-expressed genes, followed by intersection with PM I anther-upregulated genes, reduced the candidate set from 3,262 (1,035 B genes) to 7 (2 B genes). Numbers in parentheses indicate B-encoded genes. Dashed arrows indicate genes upregulated in +B. *B nondisjunction occurs in PM I anthers. (**b**) Top 10 enriched GO terms among 949 B genes upregulated in B elimination-associated tissues after primary root subtraction. Dot size indicates gene count; color indicates adjusted *P* value.

GO enrichment analysis of the 949 retained B chromosome genes revealed strong overrepresentation of terms associated with chromosome organization and cell division (Fig. 2b, Supplementary Table 4). The most significantly enriched category was nuclear chromosome segregation, followed by terms related to sister chromatid cohesion, kinetochore assembly, mitotic cell cycle progression, and centromeric region localization (Fig. 2b, Supplementary Table 4). Within the chromosome segregation category, multiple components of the centromere, kinetochore, and cohesin machinery were upregulated (Supplementary Table 5), including kinetochore components *NDC80*, *MIS12* and *NUF2*, cohesin subunit *SMC3*, and kinesin family members involved in spindle organization. Notably, this set included a B-encoded centromeric histone H3 variant (*CENH3-B*). Furthermore, transcripts typically associated with meiotic chromosome dynamics were also enriched in these mitotically dividing elimination tissues. Together, this functional enrichment supports a model in which B elimination is accompanied by coordinated upregulation of B-encoded genes targeting multiple layers of the mitotic segregation machinery.

Further subtraction of genes upregulated in +B shoots, in which Bs are stably retained, narrowed the set to 486 genes, with 12 B-encoded genes remaining (Fig. 2a, Supplementary Table 6). Although shoot expression does not necessarily exclude genes from playing a role in B elimination, this step prioritized candidates with expression patterns specifically correlated with elimination-active tissues. Final intersection with genes upregulated in PM I anthers, where B nondisjunction and drive occur, yielded a candidate set of seven genes including two B-encoded genes: *SYN2* (also referred to as *RAD21.1*, *AT5G40840*) ^44^ and an ankyrin repeat family protein (Fig. 2a, Table 2). The rationale for including PM I anthers was that B nondisjunction underlies both drive and elimination ^35^, and genes expressed during both processes are likely to represent core regulators of B chromosome missegregation. The B-encoded *SYN2* variant(*SYN2-B*), retained through to the final candidate set and centromere-specific Histone H3 (*CENH3, HTR12*) ^45^, identified during the primary root subtraction step, were selected for functional characterization based on their central roles in sister chromatid cohesion and centromere identity, respectively.

**Table 2.**
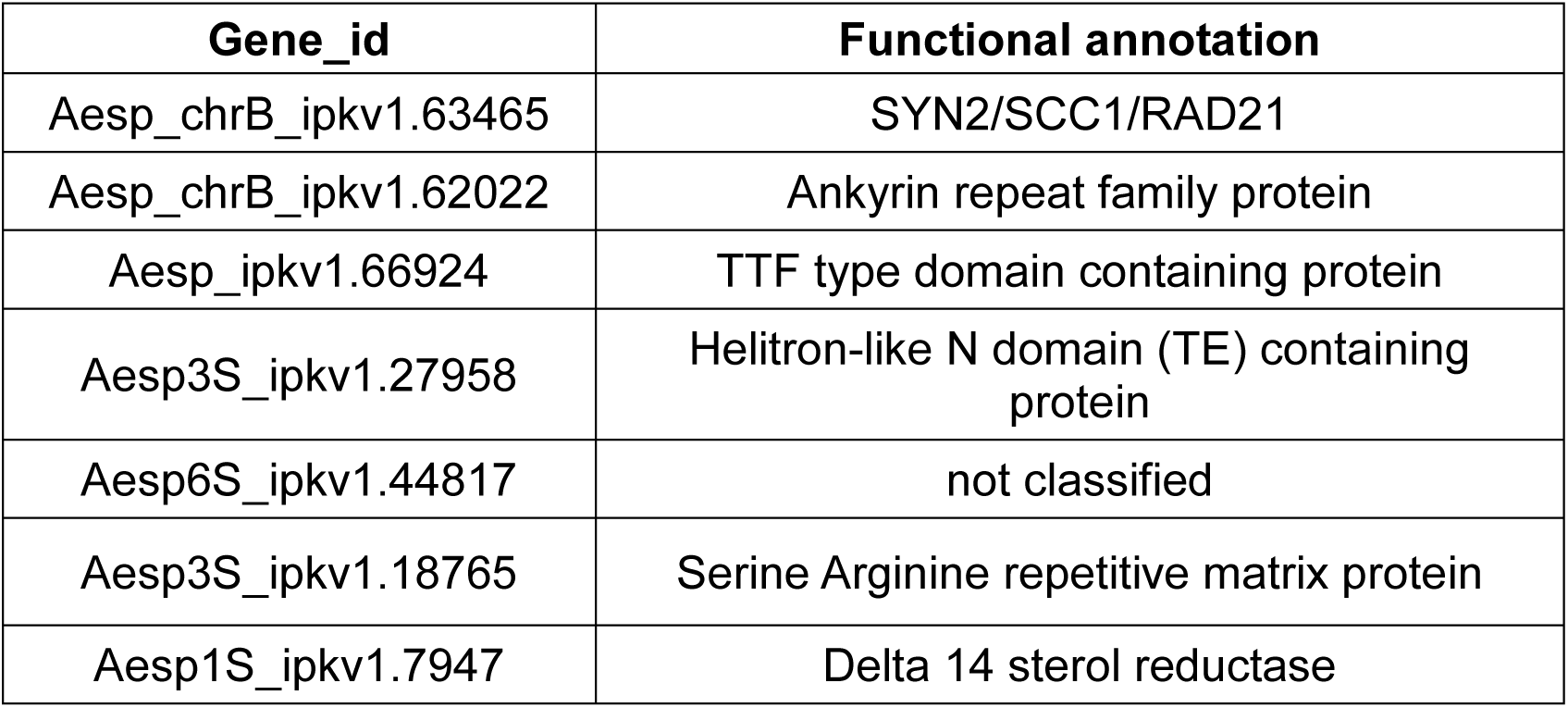
Final candidate genes identified by stepwise transcriptome-based prioritization. B genes are indicated by the prefix ‘Aesp_chrB’.

### *SYN2-B* is expressed in B chromosome nondisjunction-relevant tissues and its misregulation induces chromosome segregation defects

Among the final candidate genes, *SYN2-B* was identified as a B-encoded homolog of the α-kleisin cohesin subunit SCC1/RAD21/SYN2 ^44, 46^. Sequence analysis revealed 89% amino acid identity between SYN2-B and its A chromosome-encoded paralog SYN2-A, with strong conservation across the RAD21/REC8 cohesin domain and the central HEAT-repeat binding domain (Supplementary Fig. 7). Promoter analysis identified notable differences in predicted transcription factor binding site composition, including enrichment of the ethylene response factor (ERF) ^47^ and C2H2 family sites ^48^ for *SYN2-B*, suggesting divergent transcriptional regulation between the two paralogs (Supplementary Table 7).

Expression profiling across 7 different tissue types with and without Bs (Supplementary Table 1) revealed distinct patterns for *SYN2-A* and *SYN2-B* (Fig. 3a). *SYN2-A* exhibited constitutive expression across all tissues regardless of B presence, consistent with its role as the canonical cohesin subunit. In contrast, *SYN2-B* transcripts were detected exclusively in +B tissues undergoing B nondisjunction, including embryos and AR buds at elimination-active stages and PM I anthers, and were completely absent in all 0B samples and +B tissue not subjected to chromosome elimination (Fig. 3a). The expression pattern of SYN2-A and SYN2-B was validated by RT-PCR (Fig. 3b). In contrast, *SYN2-A* is active in all analyzed samples irrespective of B presence. Together, these results indicate that *SYN2-B* transcription is tightly coupled to developmental contexts associated with B chromosome missegregation.

**Fig. 3.**
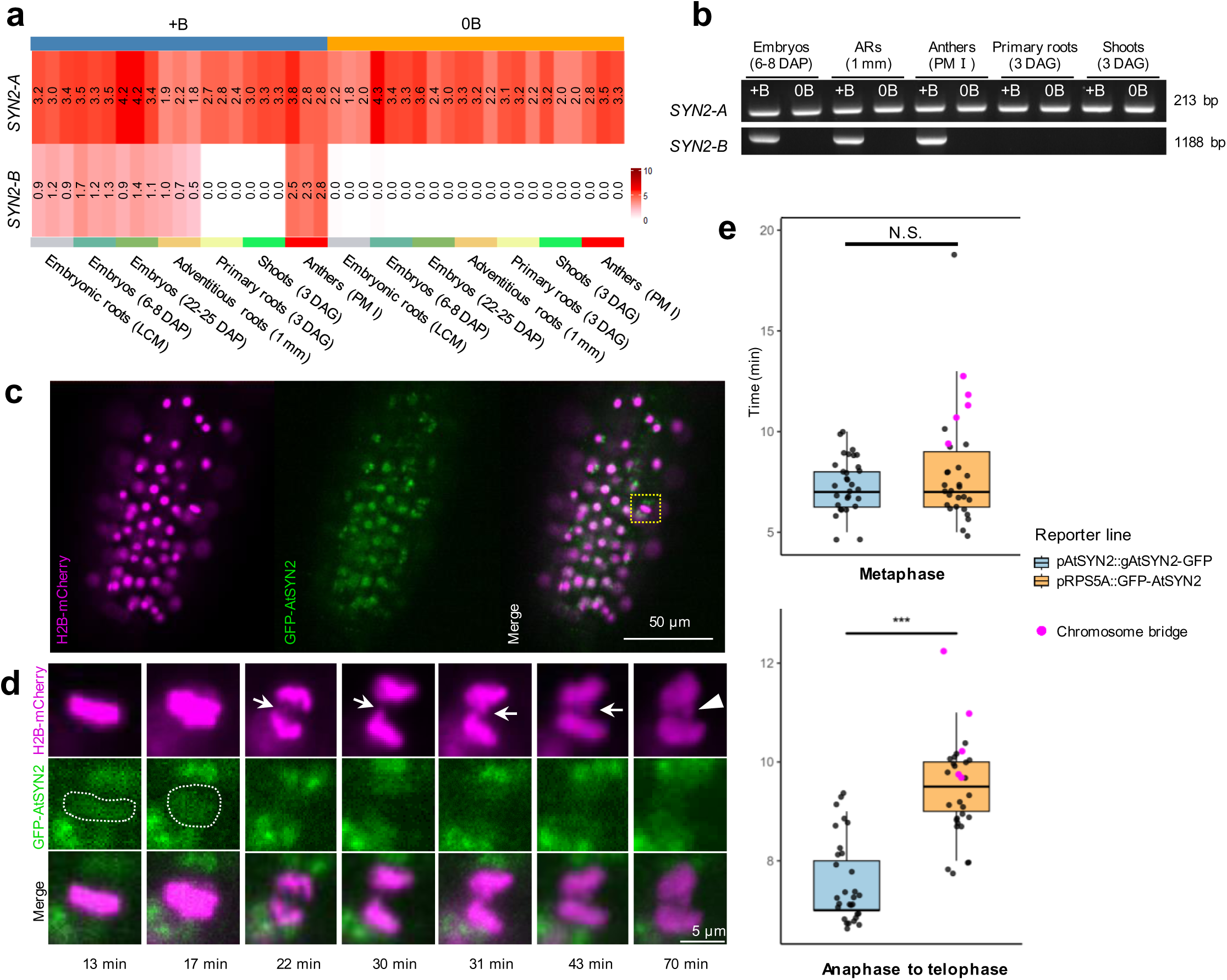
SYN2-B is expressed in +B nondisjunction-relevant tissues and its misregulation induces chromosome segregation defects. (**a**) Heatmap of *SYN2-A* and *SYN2-B* expression (log2 TPM) across 42 RNA-seq samples from 0B and +B plants. (**b**) RT-PCR validation of *SYN2-A* and *SYN2-B* expression in selected tissues. Expected amplicon sizes: *SYN2-A*, 213 bp; *SYN2-B*, 1188 bp. (**c**) Light sheet fluorescence microscopy image of *A. thaliana* root tip cells coexpressing pRPS5A::GFP-AtSYN2 (green) and histone H2B-mCherry (magenta). (**d**) Selected time frames from live-cell imaging in (c) showing chromosome bridge formation (arrow) and subsequent micronucleus formation (arrowhead). The reticulate green signal corresponds to GFP-AtSYN2 (dotted outlines, 13 and 17 min). (**e**) Duration of metaphase (top) and anaphase-to-telophase transition (bottom) in *A. thaliana* pAtSYN2::AtSYN2-GFP and pRPS5A::GFP-AtSYN2 reporter lines. Individual cells are shown as dots; magenta dots indicate bridge-forming cells. Boxplots show the median, interquartile range, and whiskers extending to 1.5× IQR. Significance was assessed using a two-sided Student’s t-test on individual cells (ten cells per line, three independent lines per construct). N.S., not significant; ***, P < 0.001.

To further investigate the potential function of the B-encoded *SYN2*, we used the model plant *Arabidopsis thaliana*, which offers more tools for functional gene analysis. *A. thaliana* encodes the single-copy orthologous gene *AtSYN2*. Previous studies showed that *AtSYN2* encodes a conserved α-kleisin subunit of the mitotic cohesin complex, where it bridges the two Structural Maintenance of Chromosomes (SMC) ATPase subunits (SMC1 and SMC3) to form the ring-like structure that holds sister chromatids together ^44, 46^. *AtSYN2* transcripts are detected throughout the plant in actively dividing tissues, with the highest levels in meristematic regions ^44^. GFP-tagged AtSYN2 localizes to chromatin from interphase through metaphase and is rapidly removed at anaphase onset, consistent with separase-mediated cleavage of the kleisin subunit that triggers sister chromatid separation ^49^.

A homozygous *syn2* knockout line (T-DNA mutant line SALK_044851) was verified by RT-PCR, confirming the absence of *AtSYN2* transcripts (Supplementary Fig. 8a). Chromosome segregation was analyzed in anaphase cells of cotyledons from 3-day-old seedlings. Chromosome bridges were observed in 14 of 133 anaphase cells (10.50%) in *syn2* mutants (31 plants), compared with 3 of 507 (0.59%) in wild-type Col-0 controls (81 plants) (Table 3, Supplementary Fig. 8b-c). Full rescue of this phenotype was achieved in three independent complementation lines expressing the genomic *AtSYN2* sequence under its native promoter (a ∼2 kb region upstream of the start codon; transgene expression confirmed by RT-PCR, Supplementary Fig. 8a). In complementation lines chromosome bridges were completely absent (0 of 144 anaphase cells, 35 plants) and indistinguishable from wild type (Table 3), thereby confirming that the segregation defects were specifically caused by loss of AtSYN2 function.

**Table 3.**
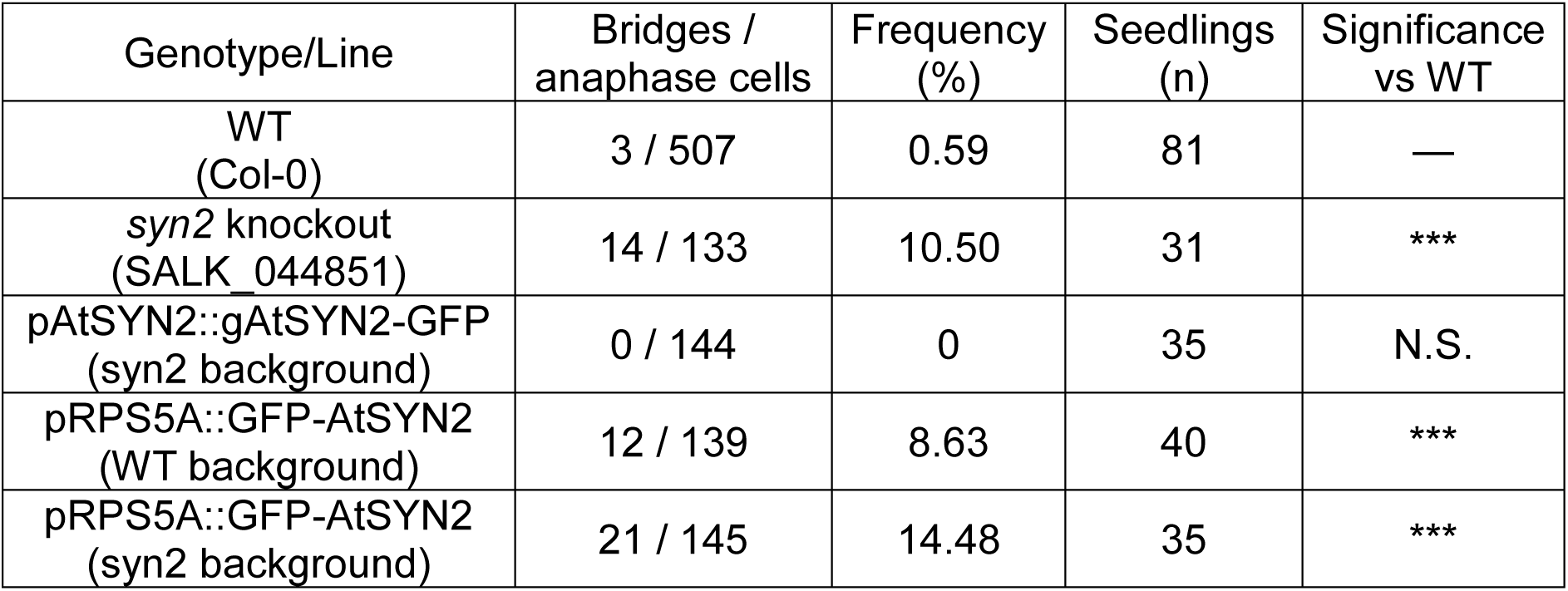
Frequency of chromosome bridges during anaphase in AtSYN2 loss-of-function, complementation, and overexpression lines. N.S., not significant, \*\*\**P* < 0.001, significant differences compared to Columbia wild type (Col-0).

Because *SYN2-B* is upregulated rather than silenced during B elimination, the effect of elevated SYN2 activity was assessed in lines expressing GFP-tagged AtSYN2 under the control of the ubiquitously-active RPS5A promoter. Chromosome bridges were detected in 12 of 139 anaphase cells (8.63%) across three independent overexpression lines in a wild-type background (40 plants) (Table 3, Supplementary Fig. 8a), demonstrating that both loss and overexpression of AtSYN2 are sufficient to compromise mitotic chromosome segregation. Expression of GFP-AtSYN2 under the RPS5A promoter in the *syn2* background further increased bridge frequency to 21 of 145 anaphase cells (14.48%) (35 plants) (Table 3, Supplementary Fig. 8a), consistent with a sensitized genetic background effect. Anti-GFP immunostaining confirmed nuclear GFP-AtSYN2 signal in root tip cells of the overexpression line (Supplementary Fig. 8d).

To directly visualize the sequence of mitotic defects, live-cell imaging of root meristems was performed in lines co-expressing GFP-AtSYN2 and the chromatin marker H2B-mCherry (Fig. 3c). In chromosome bridge-forming cells of the RPS5A promoter-driven overexpression lines, the chromosomes failed to separate regularly at anaphase onset, forming bridges that remained under tension as spindle forces acted on the connected chromatids, a behavior not observed in endogenous promoter reporter lines (Fig. 3d, Supplementary Fig. 9, Supplementary Movie 1, 2). By the end of mitosis, the lagging chromatin was excluded from both daughter nuclei and formed a micronucleus. Quantitative analysis of mitotic progression (three independent lines per construct, ten cells per line) showed that anaphase-to-telophase duration was significantly prolonged in bridge-forming cells of the overexpression lines relative to non-bridge cells and to the endogenous promoter reporter lines (Fig. 3e, Supplementary Movie 1).

Thus, both loss and overexpression of *AtSYN2* disrupt faithful chromosome segregation, resulting in chromosome bridges, lagging chromosomes, and micronucleus formation in *A. thaliana.* Given that *SYN2-B* is specifically upregulated in *Ae. speltoides* tissues where Bs undergo nondisjunction, elevated SYN2 activity is a plausible contributor to the cohesin dysregulation that promotes B-specific lagging and elimination in root tissues. Read mapping and BLAST analysis of genomic data from three additional B-carrying *Ae. speltoides* accessions (H7, D2, P12) confirmed the presence of *SYN2-B* on B-assigned scaffolds in all three genotypes, indicating that this gene is a conserved feature of the *Ae. speltoides* B chromosome.

### Structural modeling predicts that SYN2-B retains SCC3-B binding with altered interface residues

Transcriptome analysis identified a B-encoded SCC3 homolog *SCC3-B* (Aesp_chrB_ipkv1.61852, Supplementary Table 3), expressed specifically in +B tissues simultaneously with *SYN2-B*, raising the possibility that a B-encoded cohesin interaction framework operates during elimination. To investigate whether SYN2-B could modulate cohesin function within such a framework, AlphaFold3 structural predictions of SYN2-B interactions with SCC3-B were performed (Fig. 4, Supplementary Table 8, 9). While the N- and C-termini of kleisins such as SYN2 interact with the head domains of the SMC proteins, the central region interacts with HAWK proteins such as SCC3 (Fig. 4d; van Hooff, Raas ^50^). Consistent with this general kleisin–HAWK interaction pattern, the predicted interaction geometry between SYN2-B and SCC3-B is broadly conserved with that of SYN2-A, arguing against major structural incompatibility. However, several substitutions at predicted SCC3-B interface surfaces, including Cys440A/Arg441B, Asn441A/Asp442B, and Cys427A/Arg428B, favor the SYN2-B variant at key contact sites, with several of these substitutions occurring at positions otherwise conserved among SYN2 orthologs across cereals (Supplementary Fig. 10), raising the possibility of subtly altered interaction properties within a potential B-linked cohesin complex (Fig. 4e-f, Supplementary Table 8). Superposition of a predicted SYN2 central motif onto the experimentally resolved human SA2-SCC1 subcomplex (PDB: 4PJU) further supports the structural plausibility of the predicted interaction mode (Fig. 4g-h, Supplementary Table 9).

**Fig. 4.**
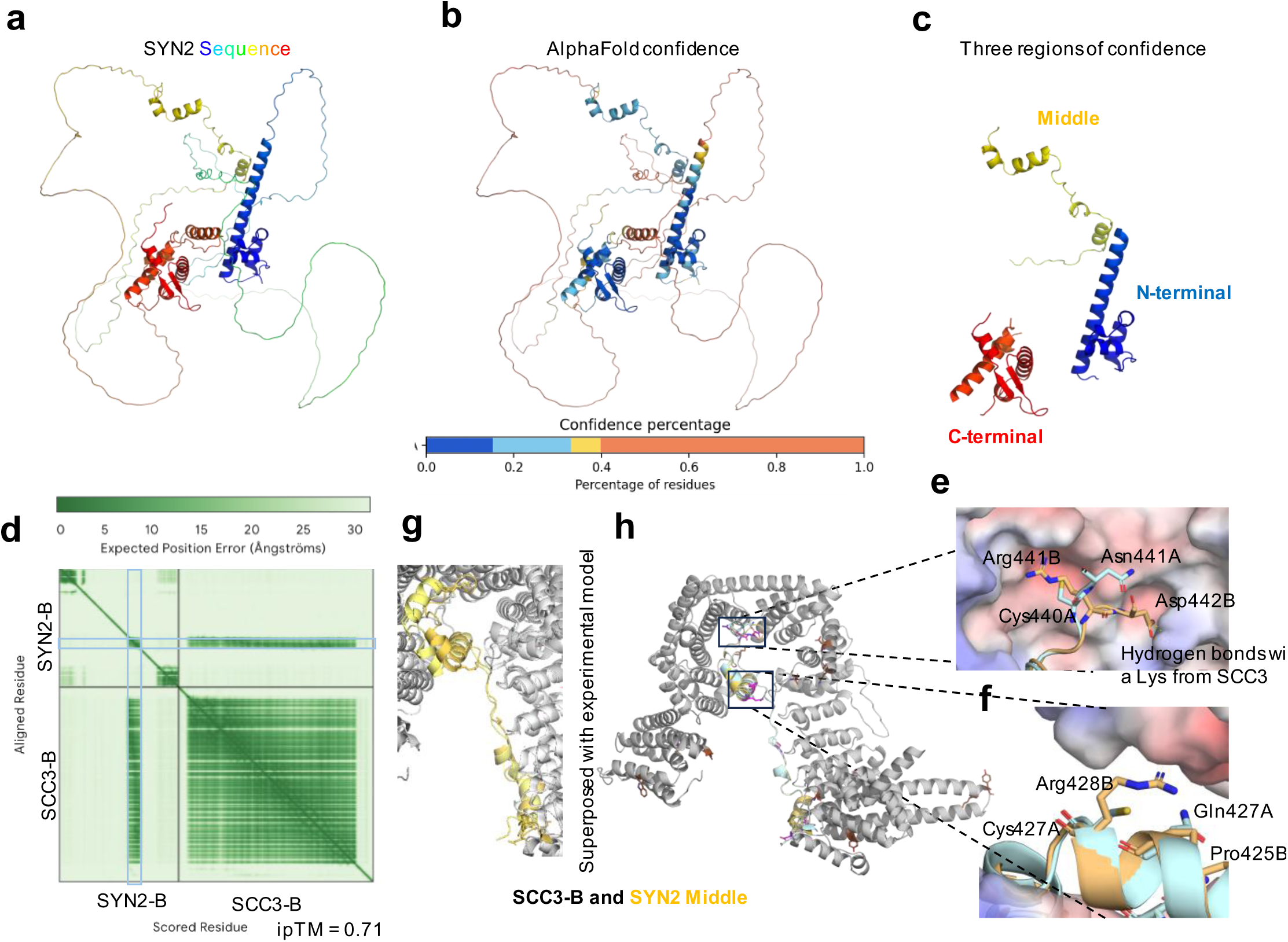
AlphaFold3 structural prediction of SYN2 interfaces with SCC3-B. (**a**) Predicted structure of SYN2-A. (**b**) AlphaFold confidence distribution for SYN2-A. (**c**) Three higher-confidence regions of SYN2-A (N-terminal, middle, and C-terminal motifs) separated by low-confidence loop regions. (**d**) Predicted alignment error map for the SYN2-B and SCC3-B interaction, including the interface predicted TM-score (ipTM). (**e–f**) Close-up views of representative SYN2-A versus SYN2-B substitutions mapped to the predicted SYN2–SCC3-B interface. Residues are labeled with A or B to indicate the corresponding SYN2 copy; residue numbering follows the respective SYN2-A or SYN2-B sequence. (**g**) Superposition of the predicted SYN2 middle-motif interaction onto an experimentally determined human SA2 (SCC3)–SCC1 (SYN2) cohesin subcomplex (PDB: 4PJU). (**h**) Superposition of the predicted SYN2-A (light blue) and SYN2-B (light orange) middle motif interacting with SCC3 (gray); amino acid substitutions of SYN2-B are colored in pink.

### A and B chromosome-encoded CENH3 variants intermingle in both chromosome types

Given that centromere identity is a key determinant of chromosome segregation fidelity, and that B chromosomes retain centromere-associated features, such as microtubule attachment, during elimination ^31^, attention has been directed toward the CENH3 protein encoded by both the A and B chromosome variants of *Ae. speltoides*.

Phylogenetic analysis of full-length CENH3 and canonical histone H3 protein sequences from *Ae. speltoides*, related Poaceae species, and other monocots revealed that the CENH3 variant encoded by the B chromosomes (CENH3-B) clusters within the αCENH3 clade, alongside the A chromosome-encoded variant (CENH3-A) of *Ae. speltoides* and centromeric histones of wheat, barley, and related grasses (Fig. 5a). No B chromosome-encoded βCENH3 paralog was identified for CENH3-B, classifying it unambiguously as an α-type variant. Sequence comparison showed strong conservation in the histone fold domain between CENH3-A and CENH3-B, with the most pronounced divergence confined to the N-terminal tail (Supplementary Fig. 11). Since all subsequent analyses addressed only α-type variants, αCENH3-A and αCENH3-B are hereafter referred to as CENH3-A and CENH3-B, respectively. *CENH3-B* transcripts were detected exclusively in +B tissues across the different sample datasets, while *CENH3-A* was constitutively expressed regardless of B chromosome presence (Supplementary Fig. 12). Similarly, *CENH3-B* was identified on B-assigned scaffolds in all three additional *Ae. speltoides* accessions (H7, D2, P12) by read mapping and BLAST analysis.

**Fig. 5.**
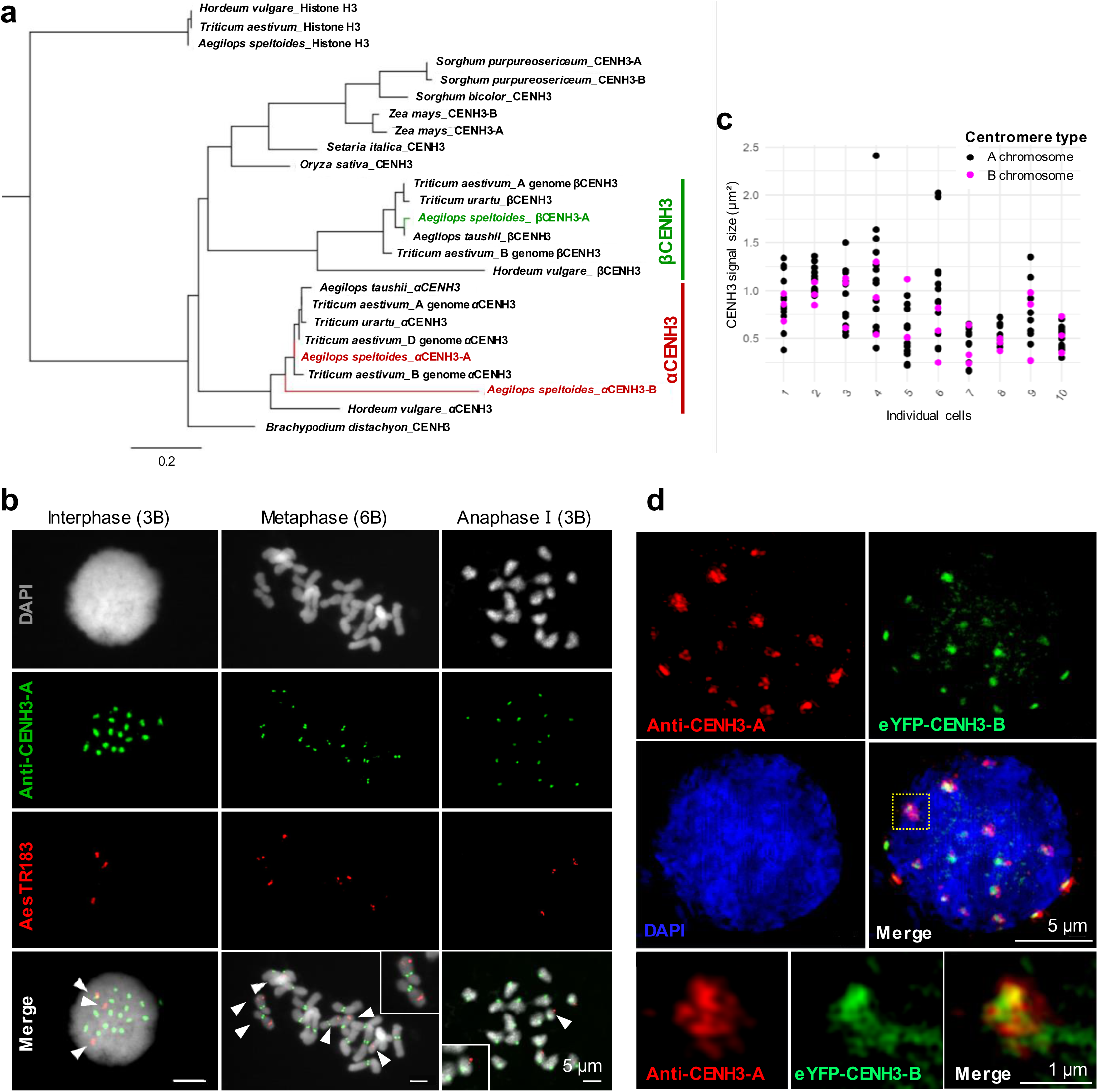
CENH3-B is an α-type centromeric histone H3 that co-incorporates with CENH3-A at centromeres of both A and B chromosomes. (**a**) Maximum likelihood phylogenetic tree of full-length CENH3 and histone H3 protein sequences from selected monocot species. *Ae. speltoides* CENH3 are highlighted. Scale bar indicates amino acid substitutions per site. (**b**) Immuno-FISH using anti-CENH3-A (green) and the B-specific probe AesTR183 (red) in interphase (3B), metaphase (6B), and anaphase I (3B) nuclei of *Ae. speltoides.* Arrowheads indicate B chromosomes. (**c**) Quantification of CENH3 signal size at meiotic metaphase I and anaphase I. Each dot represents an individual centromeric region; A chromosomes in black, B chromosomes in magenta. (**d**) 3D-SIM of a +B *Ae. speltoides* protoplast nucleus co-expressing eYFP-CENH3-B (green) and immunostained with anti-CENH3-A (red). Bottom panels show the enlarged region indicated by the dotted rectangle.

Antibodies raised against the CENH3-B N-terminal peptide did not yield reliable immunosignals. To determine whether CENH3-A occupies centromeres of both chromosome types, immuno-FISH was performed using an anti-CENH3-A antibody in combination with a B-specific FISH probe. Anti-CENH3-A signals were detected at the centromeres of both A and B chromosomes at mitotic metaphase and in meiotic cells of *Ae. speltoides*, demonstrating that CENH3-A is not restricted to A centromeres (Fig. 5b). To evaluate whether centromere activity differed between A and B chromosomes, CENH3 signal size was quantified as a proxy for centromere occupancy at meiotic metaphase I and anaphase I. No significant difference in signal size was detected between A and B centromeres, indicating comparable centromeric CENH3 levels across both chromosome types (Fig. 5c).

Although these results established that CENH3-A occupies B centromeres, whether CENH3-B itself is capable of centromeric incorporation remained unresolved. Because stable transformation of *Ae. speltoides* is not yet feasible, a transient protoplast expression system was used to assess the localization competence of both variants. CENH3-A tagged with mScarlet and CENH3-B tagged with eYFP were co-expressed under the maize ubiquitin promoter in both 0B and +B *Ae. speltoides* leaf protoplasts (Supplementary Fig. 13). Both variants localized to the nucleus and formed discrete punctate signals, and colocalized at centromeric foci in 208 of 500 protoplasts scored (41.6%). Importantly, CENH3-B centromeric localization was observed in 0B protoplasts, demonstrating that B chromosome DNA is not required for CENH3-B incorporation.

To resolve the spatial relationship between the two variants at the ultrastructural level, 3D structured illumination microscopy (3D-SIM) was performed on +B protoplast nuclei co-expressing eYFP-CENH3-B and immunostained with anti-CENH3-A. Both variants appeared as discrete dot-like chromatin-associated structures distributed throughout the nucleus, and their signals were intermingled rather than forming spatially distinct domains (Fig. 5d). Thus, the B chromosome-encoded αCENH3 variant co-incorporates together with αCENH3-A at centromeres of A and B chromosomes.

### Histone H3S10 phosphorylation dynamics do not differ between segregating and lagging A and B chromosomes

In plants, cell cycle-dependent post-translational phosphorylation of histone H3 at serine 10 is associated with sister chromatid cohesion ^51, 52^. Notably, in fungus gnats and songbirds, lagging X chromosomes and germline-restricted chromosomes (GRCs) do not undergo histone H3S10 dephosphorylation in mitotic anaphase, while all other chromosomes do ^53, 54^. To test whether the B chromosome of *Ae. speltoides* shows the same chromatin difference during elimination, we immunostained anaphase chromosomes of the proto-root tissue and identified the lagging Bs with a B-specific FISH probe. Unlike in animals, the degree of histone H3S10 phosphorylation does not differ between separated A chromatids and lagging B chromatids (Fig. 6). Thus, an epigenetic marking of the to-be-eliminated chromosome by histone phosphorylation is not a universal feature across species.

**Fig. 6.**
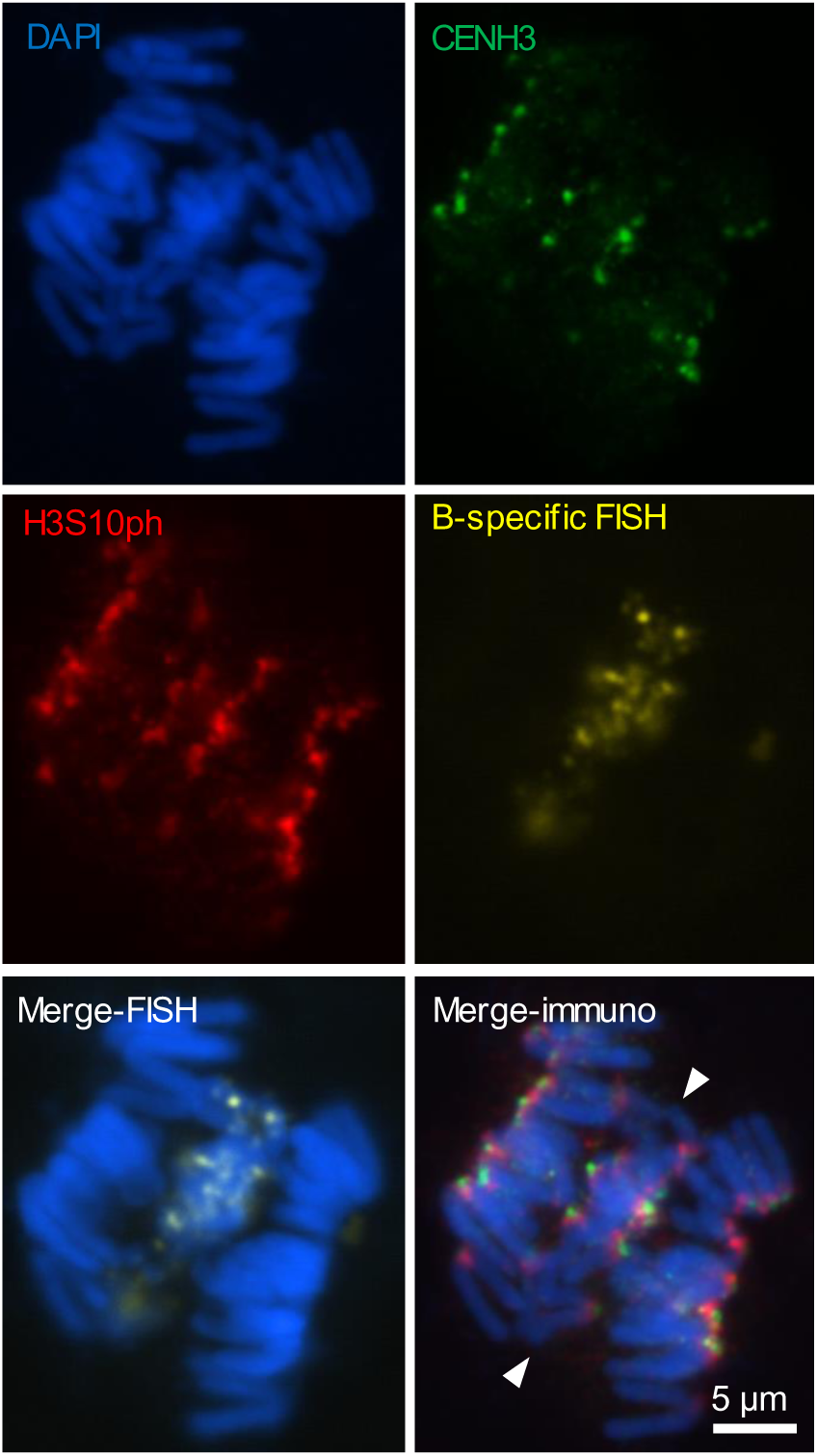
Histone H3S10 phosphorylation does not differ between segregating and lagging A and B chromosomes. Mitotic anaphase chromosomes of +B *Ae. speltoides* showing lagging B chromosomes (arrowed) after immunostaining with anti-H3S10ph (red) and anti-CENH3 (green) and FISH using the B-specific probe AesTR183.

## Discussion

This study establishes an integrated genomic and transcriptomic framework for investigating the molecular basis of programmed B chromosome elimination in *Ae. speltoides*. By exploiting the natural absence of B chromosomes in lateral adventitious and embryonic roots, a chromosome-scale genome assembly was generated from a single clonal +B plant. This high-quality assembly enabled discrimination between A-and B-encoded sequences. Consequently, it provided a robust reference for genome-resolved, tissue-specific transcriptome analysis. Multi-tissue comparative transcriptomics across B elimination-active and B elimination-negative developmental stages identified a candidate gene set enriched for functions in chromosome segregation. From this dataset, SYN2-B and CENH3-B emerged as the primary candidates for functional characterization. The remaining candidate, an ankyrin repeat family protein ^55^, was not pursued further but remains available for future investigation.

Gene Ontology enrichment analysis of B-encoded genes upregulated in elimination-associated tissues revealed strong overrepresentation of terms related to nuclear chromosome segregation, sister chromatid cohesion, kinetochore function, and centromeric region organization. This functional profile is consistent with the cytological observation that B chromosomes undergo selective nondisjunction during mitotic divisions of cells destined to become root cells, while A chromosomes segregate properly in the same division ^31^. A comparable enrichment of chromosome segregation-related GO terms among B-encoded candidates was independently identified in wild sorghum embryos undergoing B elimination ^32, 56^, suggesting that transcriptional perturbation of mitotic segregation machinery by B-encoded genes may represent an evolutionarily conserved feature of programmed chromosome elimination (Supplementary Table 10). Notably, a B chromosome-encoded SMC3 homolog and CENH3 variant were among the shared candidates identified across both species, further supporting the relevance of cohesin and centromere-related pathways in this process.

*SYN2-B* is a B-encoded cohesin α-kleisin subunit whose expression is restricted to tissues undergoing B chromosome nondisjunction, including elimination-active embryonic root zones, AR buds, and PM I anthers. In contrast, its A chromosome paralog SYN2-A is constitutively expressed. In *A. thaliana*, both loss and overexpression of the orthologous *AtSYN2* significantly increased the frequency of anaphase chromosome bridges and micronuclei, demonstrating that chromosome segregation fidelity is sensitive to SYN2 activity. Homozygous *atsyn2* knockouts remain viable, as the mitotic α-kleisins SYN2 and SYN4 can partially compensate for one another, and *syn2* mutants display anaphase bridges at frequencies close to those we observed ^46^. In canonical cohesin, the α-kleisin subunit bridges the cohesin ring and provides a direct interaction platform for the HEAT repeat subunit SCC3/SA, thereby coupling cohesion establishment and release to defined protein contacts within the complex ^57^. Mechanistic interpretation therefore relied on whether SYN2-B could plausibly influence cohesin behavior based on established cohesin architecture and structural predictions.

Together, the expression and structural analyses support a model in which *SYN2-B* upregulation in elimination-relevant tissues contributes to cohesin dysregulation, disproportionately affecting B chromosome cohesion, promoting lagging and subsequent elimination, while A chromosome segregation remains largely unaffected.

At the molecular level, HAWK proteins are increasingly recognized as active modulators of cohesin dynamics rather than passive scaffolding subunits. NIPBL and PDS5, which share the HEAT-repeat architecture of SCC3, were recently shown to tune the rate of cohesin loop extrusion and the lifetime of cohesin–DNA engagement ^58^. By analogy, a B-encoded kleisin–HAWK pair with subtly altered interface properties, such as the SYN2-B–SCC3-B module, could shift the affinity or residence time of cohesin on chromatin. Such a change might preferentially affect the duration of sister chromatid association on the B chromosome, providing a plausible mechanistic link between B-specific cohesin composition and the delayed cohesion release that accompanies B chromatid lagging.

The link between compromised sister chromatid cohesion and chromosome missegregation extends well beyond B chromosome biology. In human cells, somatic mutations in cohesin subunits and their regulators are recurrently found across diverse cancers, where they are thought to promote tumorigenesis through error-prone chromosome segregation ^59^. More generally, when cohesion is not properly regulated, cells can develop chromosomal instability, including aneuploidy, chromosome loss, and micronucleus formation, as well as cohesion fatigue ^60, 61^. This suggests that the selective, cohesion-dependent missegregation of B chromosomes is a programmed, natural equivalent to the faulty cohesion that triggers genome instability in pathological systems.

CENH3-B is a B chromosome-encoded α-type centromeric histone variant whose expression is restricted to +B tissues, including shoots (3 DAG). Phylogenetic analysis locates CENH3-B within the αCENH3 clade alongside CENH3-A and related variants of wheat and barley, indicating origin from an ancestral αCENH3 copy of *Ae. speltoides* rather than independent *de novo* evolution. Anti-CENH3-A immunostaining demonstrated that CENH3-A is present at the centromeres of both A and B chromosomes, and no significant difference in CENH3 signal size was detected between the two chromosome types, indicating comparable centromere occupancy. Transient co-expression of fluorescently tagged CENH3-A and CENH3-B in *Ae. speltoides* protoplasts, combined with 3D-structured illumination microscopy, revealed intermingled rather than spatially segregated signals, supporting a co-incorporation in which both variants can occupy centromeres of both chromosome types. B chromosome-encoded CENH3 variants have also been reported in sorghum ^32^ and maize ^62^, suggesting that acquisition of a B-specific centromeric histone may be a recurrent feature of B chromosome evolution across species. The possible function of a B chromosome-specific variant remains an open question. So far, no chromosome-type-specific loading of CENH3 variants has been reported. However, tissue-specific differences in transcription and centromere incorporation of CENH3 variants have been demonstrated ^63, 64^. For the B-specific CENH3 variant of wild sorghum, structural modeling of the CENH3–CENP-C nucleosome complex using AlphaFold3 revealed that amino acid differences between the A- and B-encoded CENH3 variants are located at the predicted interaction interface with CENP-C, with complementary changes also detected in the B-encoded CENP-C variant, suggesting the possible formation of an altered inner kinetochore on the B chromosome ^32^. Whether the combined effects of the *trans*-active SYN2-B-mediated alteration of cohesion and CENH3-B-associated centromere compositional differences are sufficient to fully account for selective B elimination, or whether additional *cis*-active B-encoded factors contribute, remains to be established through direct functional approaches in *Ae. speltoides* as transformation methods for this species become available.

The functional characterization of SYN2-B and CENH3-B supports a model in which B elimination arises from coordinated perturbations in cohesin dynamics and possibly centromere composition rather than from a single dominant molecular defect as recently discussed ^4^. B centromeres remain functionally active during elimination, as evidenced by CENH3 occupancy and microtubule attachment ^31^, yet the B chromosome selectively undergoes nondisjunction and lagging. Differences in centromere repeat composition between A and B chromosomes have been documented in *Ae. speltoides* ^35^, rye ^65^, and maize ^66^ and other species ^20^. Such differences in repeat organization may contribute to distinct centromere/kinetochore assembly properties and potentially underlie chromosome-type-specific segregation behavior during elimination. A parallel has been drawn in mammals, where chromosome-type-specific loss following centromere perturbations is influenced by centromere repeat sequence composition ^67^, suggesting that chromosome-type (peri)centromere sequence composition, or other chromosome type-specific repeats, may be a *cis*-active determinant of selective chromosome missegregation. A comparable situation has been discussed for the elimination of songbird GRCs, where centromere-associated features have been proposed to differ from standard chromosomes ^68, 69^.

How B elimination is confined to the root lineage at the level of cell fate remains an additional open question. Adventitious roots arise from a spatially restricted population of founder cells whose reprogramming toward a root fate is driven by ethylene and ERF signaling ^40^, and B elimination is initiated during a comparable developmental window in the proto-root region of embryos ^31^. ERF-mediated ethylene-related pathways have also been implicated in embryogenic transitions in *A. thaliana* ^70^. Notably, the SYN2-B promoter is enriched for ethylene response factor binding sites relative to its A-encoded paralog (Supplementary Table 7), raising the hypothesis that an ethylene-influenced root-identity program specifying these cells also activates elimination-promoting B chromosome encoded genes such as SYN2-B. According to this model, the tissue specificity of B elimination would emerge from co-opting a root developmental network, a link that could be validated by dissecting the *cis-*regulatory inputs of B-encoded candidates and by perturbing root-identity regulators.

More broadly, programmed B chromosome elimination offers a naturally occurring system in which specific chromosomes are selectively eliminated while the remaining complement segregates properly. A deeper mechanistic understanding of how such selective missegregation is encoded might provide long-term approaches for the controlled manipulation of chromosome number, a central challenge in both crop breeding and the study of aneuploidy.

## Material and Methods

### Plant material

*Aegilops speltoides* Tausch plants with or without B chromosomes, collected from Tartus, Syria (genotype K2, D2), Katzir, Israel (H7), and Ramat Hanadiv, Israel (P12) ^71^, were grown in a greenhouse under long-day conditions (16 h light at 25 °C, 8 h dark at 16 °C, 40% relative humidity, 50–60 kilolux). Plants were vernalized at 4 °C for 4 weeks at the three-leaf stage before transfer to greenhouse conditions at IPK Gatersleben (Germany). Presence of B chromosomes was confirmed by PCR using primers for the B-specific repeat AesTR-183 ^35^ (Supplementary Table 11), and exact B chromosome number was determined by chromosome counting after FISH with the same probe. Wild-type and transgenic *A. thaliana* (Columbia-0) were cultivated under long-day conditions (16 h light/8 h dark, 21 °C). Transgenic seeds were selected on ½ MS medium supplemented with cefotaxime (100 µg/ml) and either kanamycin (50 µg/ml) or hygromycin (15 µg/ml).

### Hydroponic induction of adventitious roots

A single *Ae. speltoides* plant (genotype K2) carrying three Bs was vegetatively propagated using a hydroponic AR induction system. Tillers were cut into leafy stem cuttings and placed individually in hydroponic pots with sponge supports and continuous aeration provided by an air pump. The induction solution was prepared as an iron-supplemented mixture as described in ^41^, containing 0.01 mM Fe-EDDHA (Duchefa Biochemie, Netherlands), 1.4 mM CaSO₄ (Carl Roth, Germany), and 1 mM MES (pH 5.8; Carl Roth, Germany) in distilled water. Growth conditions were: 10 h light at 22 °C (85% relative humidity) and 14 h dark at 20 °C (60% relative humidity). The development of adventitious roots was monitored daily.

### Flow cytometric analysis of genome size and B chromosome presence

To estimate genome size and B chromosome size, flow cytometric measurements were performed on leaf tissue from 0B and +B *Ae. speltoides* plants. For each measurement, 0.5 cm² of fresh leaf tissue was chopped with a sharp razor blade in a Petri dish together with an equivalent amount of Secale cereale subsp. cereale (R 737 from IPK-Gatersleben Genebank) as an internal reference standard, using either the CyStain PI Absolute P reagent kit (Sysmex-Partec, Germany) according to the manufacturer’s instructions or the nuclei isolation buffer of Galbraith, Harkins ^72^, supplemented with 1% PVP, 0.1% Triton X-100, DNase-free RNase (50 µg ml-1) and propidium iodide (50 µg ml-1). The resulting nuclei suspension was filtered through a 50 μm CellTrics filter (Sysmex-Partec, Germany), and analyzed using a CyFlow Space flow cytometer (Sysmex-Partec, Germany). Nuclei populations were identified and gated in a dotplot of fluorescence intensity versus side scatter, and the genome sizes were calculated from the G1 peak positions in the corresponding histogram. B chromosome size was estimated from the difference in genome size between the 0B plants and plants carrying three B chromosomes.

To confirm B elimination in AR tissue, nuclei were isolated from 1 mm AR buds and AR-derived lateral roots and measured as described above. The presence or absence of B chromosomes was assessed by comparing G1 peak positions with those of a 0B reference sample containing only A chromosomes. Additional peaks at higher DNA contents indicated the presence of B chromosome-containing nuclei.

### Genome sequencing, assembly and identification of B chromosome-scaffold

High-molecular-weight (HMW) DNA was extracted from leaf tissue of the clonal 3B *Ae. speltoides* plant using the NucleoBond HMW DNA kit (Macherey-Nagel, Germany). DNA quality was assessed by FEMTO Pulse (Agilent, USA) and quantified with a Quantus fluorometer (Promega, USA). HiFi library construction and sequencing using the PacBio Revio device (Pacific Biosciences, Menlo Park, CA, USA) were performed as described previously ^73^. Here, the loading concentration was 300 pM on the plate (adaptive loading), and the movie time was 30 h. The yield of one SMRT cell was 71 Gb with a mean insert length of 23.1 kb (SMRT link raw data report). All steps were conducted at IPK Gatersleben. Oxford Nanopore Technology (ONT) based libraries were generated using the SQK-LSK114 Ligation Sequencing Kit V14 (Oxford Nanopore Technologies, Oxford, UK) following the manufacturer’s protocol with 2-3 µg HMW DNA as input. R10.4.1 (FLO-PRO114M) flow cells were used for sequencing on the PromethION24 platform with a run-time of 96 hours per run. Raw data in the .pod5 format was acquired using MinKNOW (version 24.11.11). Basecalling was performed with Dorado (version 0.9.1) using the dna_r10.4.1_e8.2_400bps_sup@v4.2.0 super-accurate basecalling model with the “--min-qscore 20” parameter to yield basecalled reads in the .bam format. Reads were then converted to the .fastq.gz format using SAMtools ^74^ and filtered for a minimum length of 25 kb using SeqKit ^75^, yielding a total of 36.4 Gb ONT reads across four flow cells.

For B-depleted (0B) short-read data, genomic DNA was extracted from AR-derived lateral roots of the same 3B plant and sequenced on the DNBSEQ platform (PE150; BGI Genomics, Hong Kong), yielding 59.07 Gb of paired-end 150 bp reads. To assess conservation of B-encoded candidate genes across accessions, leaf genomic DNA from B chromosome-carrying individuals of three additional *Ae. speltoides* genotypes (H7, D2, P12) was likewise sequenced on the DNBSEQ platform (∼30 Gb per sample); read mapping and BLAST analysis against the K2 +B reference were used to confirm the presence of *SYN2-B* and *CENH3-B* on B-assigned scaffolds in all three genotypes. Hi-C chromatin interaction libraries were prepared from leaf tissue and were sequenced (paired-end, 2× 111 cycles) using the NovaSeq6000 device (Illumina Inc., USA) at IPK Gatersleben. A total of ∼104 Gb paired-end reads were generated.

Whole-genome assembly of *Ae. speltoides* carrying three B chromosomes was performed using hifiasm (v0.19.3-r572; Ultra-long ONT integration) ^42^, incorporating 154 Gb PacBio HiFi reads and 36.4 Gb ONT reads (Q20, >25 kb). Contig statistics were calculated with Quast (v2.3) ^76^ and completeness was assessed using BUSCO (v5) ^77^ against the Poaceae_odb10 lineage dataset ^78^. The Arima Genomics mapping pipeline (https://github.com/ArimaGenomics/mapping_pipeline) was used to process the Hi-C data, including read mapping to the contigs, read filtering, read pairing, and PCR duplicate removal. Scaffolding was performed using YaHS ^79^ (v1.2a.2). Hi-C contact maps were generated using Juicebox (https://github.com/aidenlab/Juicebox). MUMmer4 ^80^ was used to align the scaffolds to the publicly available *Ae. speltoides* genome ‘AEG-9674-1’ ^43^ and the results were visualized via NGenomeSyn ^81^ using the ‘GetTwoGenomeSyn.pl’ script.

To determine which scaffold belongs to the B chromosome, ∼60 Gb of whole-genome sequencing (WGS) data from 0B lateral adventitious root (AR) tissue of the same plant and 36 Gb of WGS data from +B leaf tissue were aligned to the scaffolds using bowtie2 (v2.5.0, default) ^82^. To assess the presence of the B chromosome, read coverage was normalized using *bamCoverage* from deepTools (v3.5.1) ^83^ with a bin size of 100 kb. The resulting coverage profiles were visualized by pyGenomeTracks (v3.8) ^84^.

### Genome-guided transcriptome annotation and RNA-seq analysis

For transcriptome profiling, RNA was isolated from seven tissue types of *Ae. speltoides* (genotype K2): whole embryos at 6–8 DAP and 22–25 DAP; laser-capture microdissected (LCM) embryonic roots from 17–20 DAP embryos; 1 mm AR buds; primary roots (3 DAG); shoots (3 DAG); and PM I anthers. For each tissue, 0B and +B plants were sampled in three biological replicates (42 libraries total). Embryos at defined stages were obtained by manual pollination after emasculation, with the day of anthesis recorded. All tissues were immediately frozen in liquid nitrogen and stored at −80 °C.

For LCM-based RNA isolation, 17–20 DAP embryos were cryosectioned with 16 µm thickness using a CryoStar NX70 cryotome (Thermo Fisher Scientific, USA) and mounted on PEN membrane slides (MMI, Germany). Embryonic roots were microdissected using an MMI Laser Cell Cut system coupled to an Olympus IX81 microscope. RNA was extracted using the Absolutely RNA Nanoprep Kit (Agilent, USA) and amplified by one round of T7-based *in vitro* transcription (MessageAmp II, Invitrogen) following ^36, 85^. For all other tissues, RNA was extracted using the Absolutely RNA Microprep Kit (Agilent, USA) and quality-assessed by Agilent 2100 Bioanalyzer. Sequencing libraries were prepared with the Illumina TruSeq RNA Kit v2 (Novogene UK) and sequenced on the Illumina HiSeq 2500 platform (150 bp paired-end). Raw reads were quality-trimmed with fastp v0.24.2 ^86^ (parameters: -q 30 -w 4 -l 50).

A genome-guided transcriptome annotation was generated by aligning the 21 +B libraries to the assembly using HiSAT2 ^87^ and assembling transcript models with StringTie ^88^, producing a GTF annotation comprising genes located on B-assigned scaffolds (hereafter B-encoded genes). TransDecoder (https://github.com/sghignone/TransDecoder, v5.5.0) was used to annotate coding regions within transcripts. The transcripts on the B chromosome were annotated by alignment to the protein sequences of *Ae. speltoides* genome ‘AEG-9674-1’ ^43^ via BLASTX (v 2.14.1+, e-value < 0.00001).

RNA-seq data were aligned to the *Ae. speltoides* +B genome assembly using STAR v2.7.9a ^89^ in transcriptome-guided mode (--quantMode TranscriptomeSAM), retaining uniquely mapped reads (--outSAMmapqUnique 60; --outFilterMismatchNoverReadLmax 0.04). Transcript-level quantification was performed with RSEM v1.3.3 ^90^, yielding TPM and expected count estimates.

Gene-level expected counts were imported using tximport ^91^ and analyzed separately for each tissue with edgeR ^92^. After filtering low-expression genes (filterByExpr) and TMM normalization, dispersions were estimated, and a generalized linear model was fitted; differential expression between +B and 0B samples was assessed by likelihood ratio test, with 0B samples set as the reference (baseline) group so that log2 fold changes represent expression in +B relative to 0B samples. Genes with FDR < 0.01 and |log2 fold change| > 1 were considered differentially expressed.

### Gene Ontology enrichment analysis

Functional annotation of predicted proteins was performed using eggNOG-mapper v2.1 ^93^ against the eggNOG database. Gene Ontology (GO) enrichment analysis was performed with the clusterProfiler R package ^94, 95^ using the enricher function (*P*-value cutoff 0.05; q-value cutoff 0.2; Benjamini–Hochberg multiple testing correction). The background gene set was restricted to genes with TPM ≥ 0.1 in the analyzed tissues. GO terms were obtained from GO.db (v3.21); obsolete or descriptor-lacking terms were excluded.

### Promoter analysis

Putative promoter sequences (2 kb upstream of the transcription start site) of *SYN2-A* and *SYN2-B* were extracted from the genome assembly. Transcription factor binding site (TFBS) prediction was performed using the PlantTFDB v5.0 ^96^.

### Phylogenetic analysis

CENH3 protein sequences from *Ae. speltoides* and related Poaceae species were identified by BLASTP against the PANTHER database ^97^. Sequences were aligned with MAFFT (L-INS-i algorithm) ^98^; non-homologous N-terminal regions were trimmed manually in SeaView ^99^. Maximum-likelihood phylogenetic trees were inferred with PhyML 3.1^100^ under the LG substitution model with gamma-distributed rate heterogeneity (four categories) ^101^. Branch support was assessed by SH-like approximate likelihood ratio test (aLRT) ^102^. Trees were visualized in FigTree v1.4.1 (http://tree.bio.ed.ac.uk/software/figtree/). Canonical histone H3 sequences were included as an outgroup.

### Structural prediction and analyses

Structural predictions were performed using the AlphaFold3 online server ^103^. Model quality and confidence were evaluated using the predicted local distance difference test (pLDDT) and the interface predicted template modeling (ipTM) score. The predictions were compared with the crystallographically resolved interaction between human SA2 (SCC3) and SCC1 (SYN2) (PDB: 4PJU), yielding a root mean square deviation (RMSD) of 2.477 Å over 44 SYN2 residues at the middle motif after superposition in PyMOL (Schrödinger, LLC). PyMOL was used for visual inspection, structural superposition, surface charge distribution analysis with APBS ^104^, and figure rendering. Sequence alignment was performed with MAFFT ^105^.

### Chromosome spread preparation and fluorescence *in situ* hybridization (FISH)

Mitotic and meiotic chromosome spreads were prepared from young *Ae. speltoides* inflorescences, and meiotic anthers from young spikes collected after 24 h ice-cold water treatment. Tissues were fixed in ethanol:acetic acid (3:1) for 3–6 days at 4 °C, transferred to 70% ethanol, and stored at −20 °C. Spreads were prepared by squashing meristematic tissue in 45% acetic acid, flash-frozen in liquid nitrogen, and stored in 99.8% ethanol at −20 °C.

For FISH, probes were generated from plasmid DNA encoding the B-specific repeat AesTR-183 ^35^ and the A chromosome pericentromeric repeat pBs301 ^106^, labeled with dUTP-ATTO-488 or dUTP-ATTO-550 by nick-translation (ATTO NT Labeling Kit; Jena Bioscience, Germany). Chromosomal DNA was denatured in 2 N NaOH in 70% ethanol (6 mg/ml) for 5 min at room temperature, dehydrated in a graded ethanol series, and hybridized overnight at 37 °C with 1 µl probe (50 ng/µl) in hybridization mixture (50% formamide, 10% dextran sulfate, 2× SSC, sonicated fish sperm DNA). Post-hybridization washes were performed in 2× SSC (2 × 5 min) at room temperature. Slides were mounted in 4′,6-diamidino-2-phenylindole (DAPI) antifade solution (1 µg/ml) ^107^.

For FISH on AR tissue sections, fresh AR buds (∼1 mm) were fixed using superglue (UHU instant glue, UHU Holding GmbH, Germany) on specimen discs and sectioned at 50 µm using a Leica VT1000S vibrating microtome (Leica Biosystems, Germany). Sections were fixed in ethanol:acetic acid (3:1) for 2–3 days at room temperature. FISH was performed in 0.5 ml reaction tubes: sections were denatured in 2 N NaOH/70% ethanol (6 mg/ml) for 5 min, washed through a graded ethanol series, and hybridized at 37 °C for 24 h with 50 µl hybridization mixture. Post-hybridization of tissue sections were performed in distilled water (5 min), followed by incubation in DAPI antifade solution (1 µg/ml, 10 min). Sections were placed on microscope slides and sealed with coverslips.

### Antibody generation, immunostaining, and immuno-FISH

Polyclonal rabbit antibodies were raised against synthetic peptides corresponding to the N-terminal tails of αCENH3-A (RRQETDGAGTSATPRRA-C) and αCENH3-B (LKMTRTKQAAVSKLKV-C; LifeTein, USA). For the detection of phosphorylated histone H3S10, a mouse anti-H3S10ph (Abcam, UK, cat. no. ab14955) was used. Chromosome slides were blocked with 4% BSA, 0.1% Tween-20 in 1× PBS for 30–60 min at 37 °C, then incubated overnight at 4 °C with primary antibody (1:1000 in 1% BSA/PBS). Secondary antibody (anti-rabbit Cy3, cat. 111-165-003, or anti-rabbit Alexa 488, cat. 711-545-152, or anti-mouse Alexa488, cat. no. 715-546-151; all Jackson ImmunoResearch, UK; 1:200) was applied for 1 h at 37 °C. Slides were washed three times in 1× PBS and dehydrated through a graded ethanol series before mounting in DAPI antifade solution. For immuno-FISH, immunostained slides were post-fixed in ethanol:acetic acid (3:1) for 24 h, denatured in 0.2 N NaOH for 5 min, washed in ice-cold 1× PBS, dehydrated, and subjected to FISH as described above.

### Transient expression in *Ae. speltoides* protoplasts

Protoplasts were isolated from etiolated ten-day-old *Ae. speltoides* (K2) seedlings grown in the dark at 23 °C according to Shan, Wang ^108^ and Gerasimova, Korotkova ^109^. mScarlet-αCENH3-A and eYFP-αCENH3-B fusion constructs were expressed under the maize *Polyubiquitin 1* promoter (including 5′-UTR with intron 1) in the binary vector pICSL4723. PEG-mediated transformation was performed using 5 × 10⁵ protoplasts, 20 µg plasmid DNA, in a total volume of 220 µl. Protoplasts were incubated for 48 h at 21 °C in the dark before imaging. For immunostaining during transient expression, protoplasts were harvested by centrifugation (1,000 × g, 3 min), fixed in 4% paraformaldehyde/PBS on ice (20 min), washed twice in 1× PBS, cytospun onto slides (Shandon Cytospin 3; 400 rpm, 5 min), and immunolabeled as described above.

### *A. thaliana* transformation and analysis of chromosome lagging

Coding sequences (CDS) of *AtSYN2* were cloned by Gibson Assembly into the pK7WGF binary vector under the RPS5A promoter, fused to N-terminal eGFP, with CaMV 35S terminator. A genomic *AtSYN2* construct driven by its native promoter, fused to C-terminal GFP in pGWB501 ^49^, was provided by Arp Schnittger (University of Hamburg). A 35S::H2B-mCherry cassette was co-introduced for chromatin visualization in localization experiments. Constructs were introduced into *Agrobacterium tumefaciens* strain AGL1 by electroporation and transformed into *A. thaliana* Col-0 by floral dip ^110^. Genomic PCR and RT-PCR genotyped the homozygous *syn2* T-DNA insertion line (SALK_044851) using specific primer pairs (Supplementary Table 11) to confirm the absence of *AtSYN2* transcript.

Chromosome bridge frequency was assessed in cotyledon cells of 3-day-old seedlings. Seedlings were fixed in ethanol:acetic acid (3:1) for 3 days, washed in 2× SSC, and enzymatically digested (2% pectinase, 2% cellulase in PBS; 2 h, 37 °C). Cotyledons were squashed in DAPI antifade solution and imaged with a 60×/1.4 Plan-Apochromat objective. Anaphase cells were scored for chromosome bridges and lagging chromosomes.

### Microscopy and image analysis

Widefield fluorescence images of FISH and immunostained chromosome spreads were acquired on an Olympus BX61 epifluorescence microscope equipped with an ORCA-ER CCD camera (Hamamatsu, Japan) and a deconvolution system, and processed with cellSens Dimension v1.11 (Olympus). FISH images of tissue sections were acquired on a Zeiss LSM 780 confocal laser scanning microscope with a 40×/1.2 water objective (ZENBlack v3.2). Confocal images of *Ae. speltoides* protoplasts and *A. thaliana* root tips were acquired on a Zeiss LSM 980 with a 40×/1.2 water objective (ZENBlack v3.6).

For live-cell imaging, root tips of *A. thaliana* seedlings co-expressing GFP-AtSYN2 (pRPS5A or pAtSYN2 promoter) and H2B-mCherry were imaged by light-sheet fluorescence microscopy (LSFM) using a Lightsheet 7 (Carl Zeiss GmbH) equipped with two pco.edge 4.2 sCMOS cameras (PCO AG). Seedlings were mounted into ∼2.15 mm (inner diameter) glass capillaries (size 4, blue mark; Carl Zeiss GmbH, 701910) containing 1% low-melting agarose (Sigma-Aldrich, A9045) in water. The temperature in the chamber was set to 21 °C, and no artificial source of light was supplemented. Imaging was done with 20x (W Plan-Apochromat 20x/1.0) detection and 10x (LSFM 10x/0.2 foc) illumination objectives and 1.0x zoom. Excitation was done with 10% 488 and 5% 561 nm laser lines. Dual-side illumination was chosen, and the light sheet pivot was turned on to reduce shadowing. Z-stacks covering the whole root were taken every 1 min. Images were acquired with ZEN Black 3.1 and processed (sample drift correction, partial movie fusion, movie and image export) with ZEN Blue 3.4 (both from Carl Zeiss GmbH).

Anaphase-to-telophase duration was measured from the first frame of sister chromatid separation to the last frame before nuclear envelope reformation, as marked by H2B-mCherry signal redistribution.

Super-resolution imaging was performed using three-dimensional structured illumination microscopy (3D-SIM) with a 63×/1.4 Plan-Apochromat oil objective on an Elyra 7 microscope system (Carl Zeiss GmbH) with the ZENBlack software. Image stacks were captured separately for each fluorochrome using 405, 488, and 561 nm laser lines with appropriate emission filters. SIM reconstruction was performed with the ZENBlack SIM processing module using default parameters ^111^. CENH3-A signal volumes at meiotic metaphase I and anaphase I centromeres were quantified via surface rendering using the Imaris v9.7 (Bitplane) software on 3D-SIM image stacks to determine the CENH3-A signal volumes of A and B chromosomes. All images were pseudo-colored using the respective acquisition software.

### Statistical analyses

Anaphase bridge frequencies were compared between genotypes at the level of individual anaphases using two-sided Fisher’s exact tests, as described previously ^46^. Exact 95% binomial confidence intervals for bridge frequencies were calculated using the mid-p method ^112^. Mitotic timing (metaphase and anaphase-to-telophase duration) was compared using two-sided Student’s t-tests on individual cell measurements (ten cells per line from three independent lines per construct), with bridge-forming and non-bridge-forming cells analyzed separately. *P* values are reported in the corresponding figure legends and tables.

## Data availability

All sequencing data generated in this study have been deposited in the European Nucleotide Archive (ENA). Genome assembly sequencing (PacBio HiFi, Oxford Nanopore Technologies, Hi-C), B-depleted short-read (DNBSEQ), RNA-seq data are available under project ID PRJEB106942 (Supplementary Table 12). Short-read genomic data from B-carrying individuals of four *Ae. speltoides* genotypes (K2, H7, D2, P12) used for cross-accession conservation analysis are available under project ID PRJEB89395 (Supplementary Table 12). The genome assembly and annotation of *Ae. speltoides* containing B chromosomes have been deposited in Zenodo (10.5281/zenodo.21287216).

## Acknowledgements

The authors thank Oda Weiss, Katrin Kumke, Manuela Knauft, Ingrid Otto, Sylvia Swetik, Nicole Schäfer, and Andrea Kunze (IPK) for technical help, and Konstantina Kleftogianni (IPK) for propagation of clonal *Aegilops* plants. We thank the IPK for providing the technical infrastructure. We thank Prof. Arp Schnittger (University of Hamburg) for providing the p*AtSYN2*::*AtSYN2*-GFP plasmid. A.Ho., J.T. and J.S. were supported by grants from the DFG (HO1779/34-1, TH 1876/6-1, SZ 397/3-1). A.Ho. was supported by DAAD-funded project 57601714. M.K. and T.B. were supported by Ministry of Education, Youth and Sports of the Czech Republic (project no. LUC24010). J.B. was supported by the project TowArds Next GENeration Crops, reg. no. CZ.02.01.01/00/22_008/0004581 of the ERDF Program Johannes Amos Comenius. Computational resources were provided by the e-INFRA CZ project (ID:90254), supported by the Ministry of Education, Youth and Sports of the Czech Republic.

## Contributions

G.K. performed the majority of the experiments, sequence and image analyses, including plant cross, hydroponic propagation, DNA and RNA isolation, transcriptome analysis, cloning, immunostaining, FISH, tissue sections, LCM, protoplast transient expression, *A. thaliana* transformation and chromosome segregation analysis, LSFM; J.C. genome assembly, gene annotation; J.T. conducted LCM and RNA isolation and mRNA amplification; D.L. immunostaining; T.B. *Sorghum purpureosericeum* data analysis; J.F. conducted flow cytometry; M.C. performed LSFM, movie processing and image analysis; V.S. performed super-resolution microscopy and image analysis; J.S. transcriptome analysis; A.S.C. protein structure prediction; J.B. and M.K. comparative transcriptome analysis; J.K. supervised protoplast transient expression; M.R.H. designed hydroponics system; B.P.P. cloning; S.J. performed ONT library preparation and sequencing; A.Hi. performed PacBio and Hi-C library preparation and sequencing; A.F. sequencing datasets submission; A.Ho. supervised the research project; G.K., J.C., and A.Ho. wrote the manuscript with the input from all coauthors.

## Supplementary Figures

**Supplementary Fig. 1.**
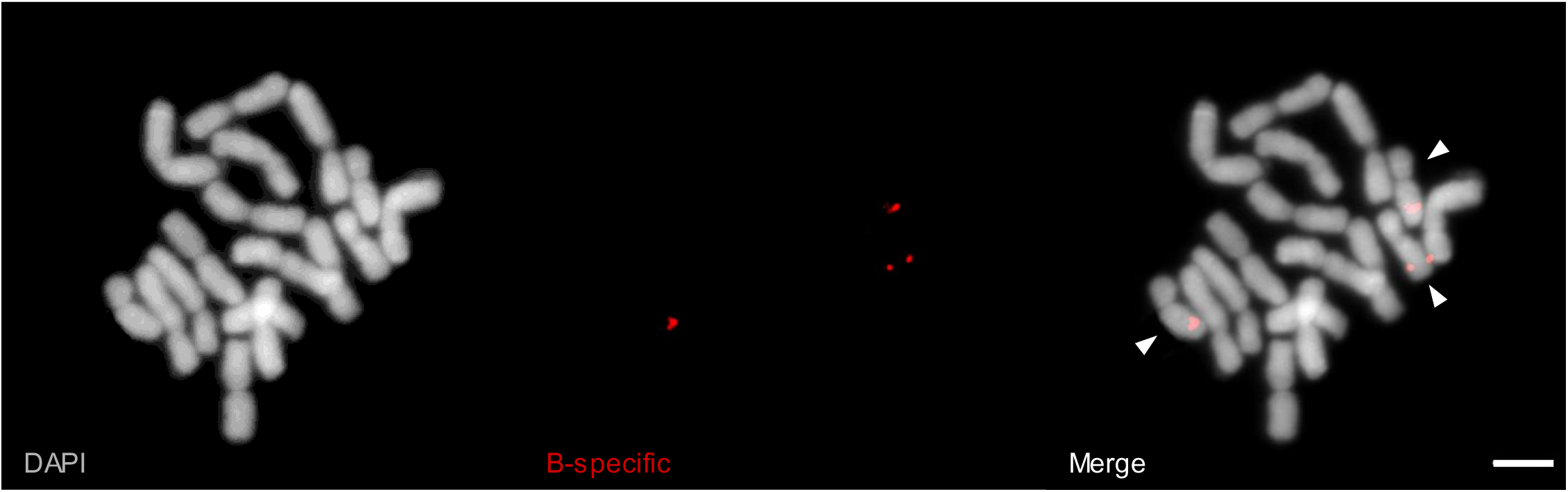
Cytological identification of B chromosomes in *Ae. speltoides* (K2, Tartus). Mitotic metaphase cell from the clonally propagated 3B *Ae. speltoides* plant used for genome assembly. FISH was performed using the B chromosome-specific repeat probe AesTR183 (red). Arrowheads indicate the B chromosomes. Chromosomes were counterstained with DAPI (gray). Scale bar: 5 μm.

**Supplementary Fig. 2.**
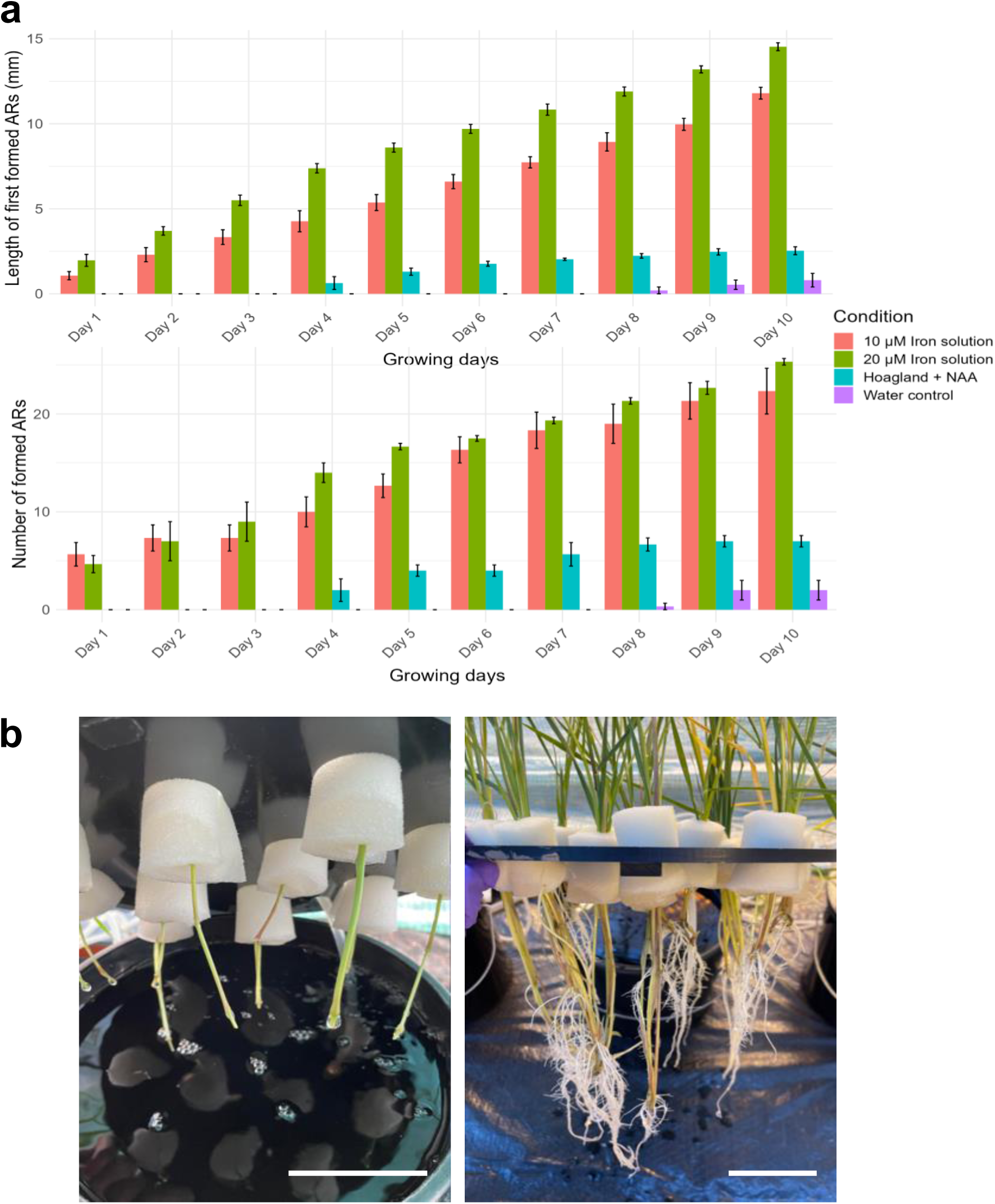
Hydroponic adventitious root (AR) induction system for clonal propagation of *Ae. speltoides*. (**a**) Mean length of the first-formed ARs (top) and mean number of ARs formed per stem cutting (bottom) over 10 days under four hydroponic conditions: 0.01 mM Fe-EDDHA solution (1.4 mM CaSO₄, 1 mM MES, pH 5.8), 0.02 mM Fe-EDDHA solution (1.4 mM CaSO₄, 1 mM MES, pH 5.8), Hoagland solution + 1-naphthaleneacetic acid (NAA), and water control. Ten cuttings were tested per condition across three independent experiments. Error bars indicate standard error of the mean. (**b**) Representative *Ae. speltoides* stem cuttings at the start of AR induction (left) and after 10 days in 0.01 mM Fe-EDDHA solution (right). Scale bar: 3 cm.

**Supplementary Fig. 3.**
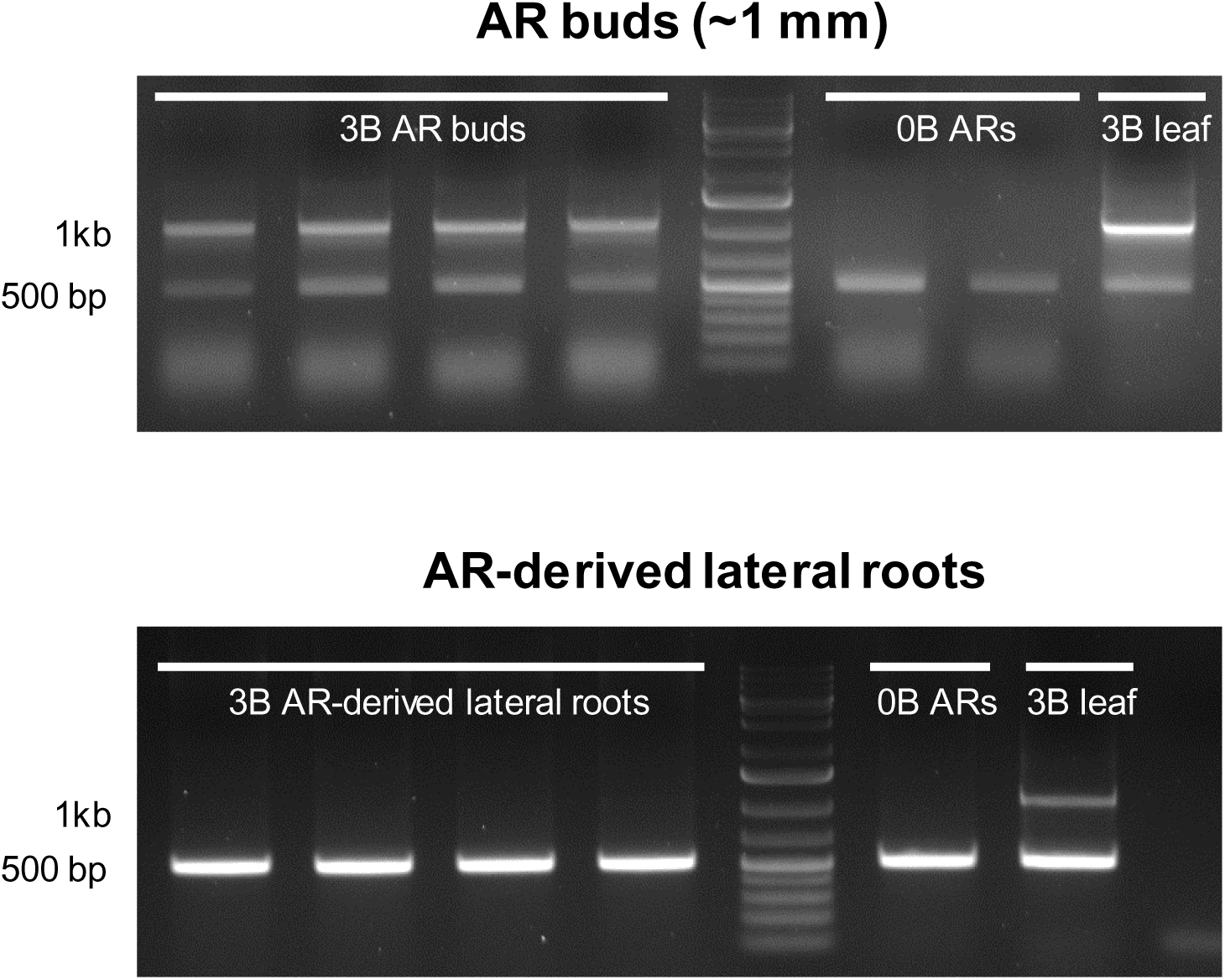
PCR-based confirmation of B chromosome elimination in adventitious root tissues. Multiplex PCR was performed on genomic DNA isolated from AR buds (∼1 mm) and AR-derived lateral roots of the clonal 3B *Ae. speltoides* plant, using a B chromosome tandem repeat-specific primer pair AesTR183 (expected amplicon size: ∼1 kb) and a *Triticum aestivum* 18S rDNA primer pair (expected amplicon size: ∼500 bp) as an A chromosome DNA control. Genomic DNA from 0B AR tissue and 3B leaf tissue were included as negative and positive controls for B chromosome presence, respectively. The B-specific band is detectable in AR buds but absent in AR-derived lateral roots, confirming that B chromosome elimination is complete in lateral root tissue.

**Supplementary Fig. 4.**
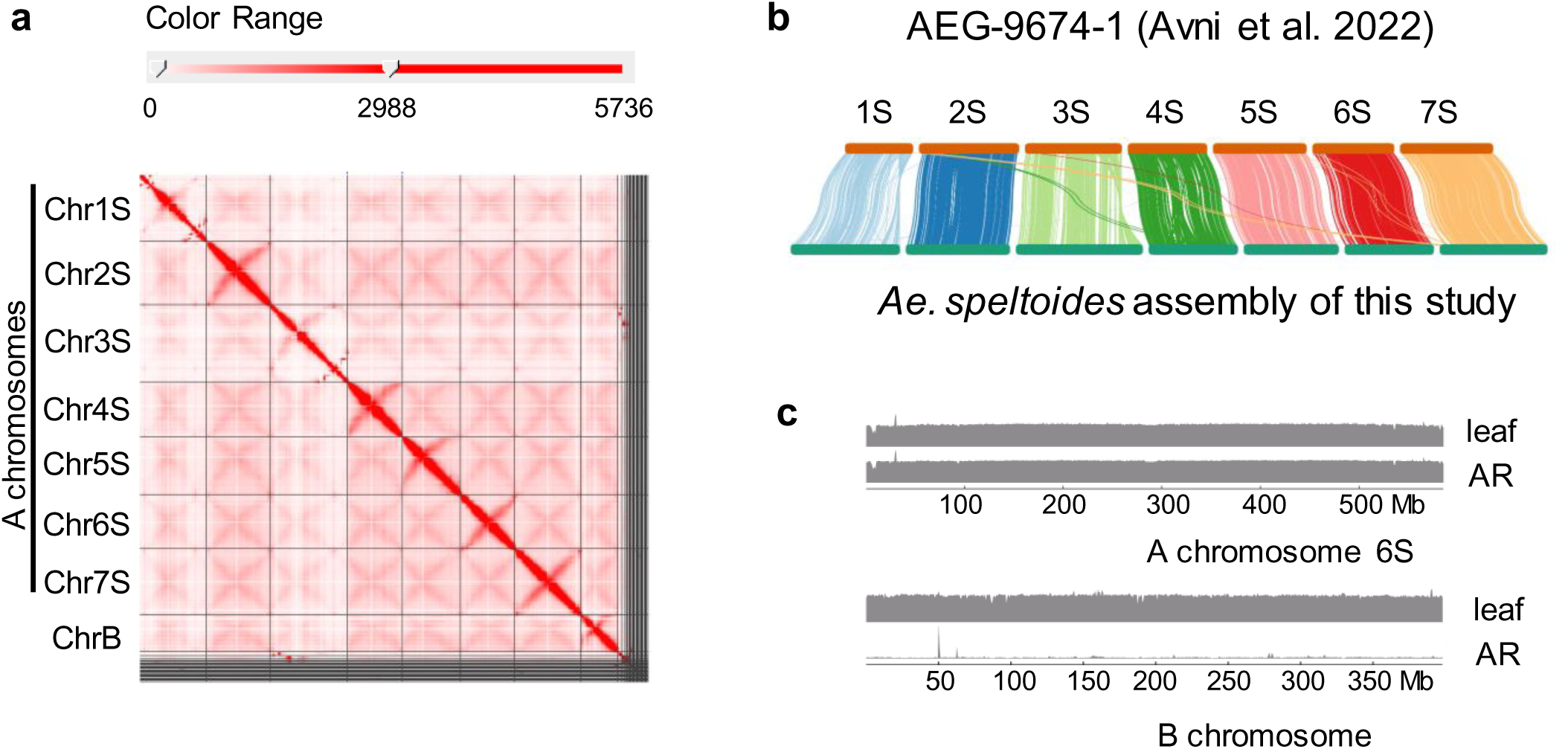
Genome assembly of *Ae. speltoides* carrying B chromosomes. (**a**) Hi-C interaction heatmap of the genome assembly of *Ae. speltoides* carrying B chromosomes. The color bar on the top represents the density of Hi-C interactions, which are indicated by the number of links at the 5-Mb resolution. (**b**) Genome alignment between the B chromosome–free reference genome of *Ae. speltoides* ^43^ and the seven longest scaffolds assembled in this study. (**c**) WGS mapping analyses showed that, unlike the A chromosomes, the assembled B chromosome exhibited normal sequencing coverage in leaf-derived data but substantially reduced coverage in AR-derived data.

**Supplementary Fig. 5.**
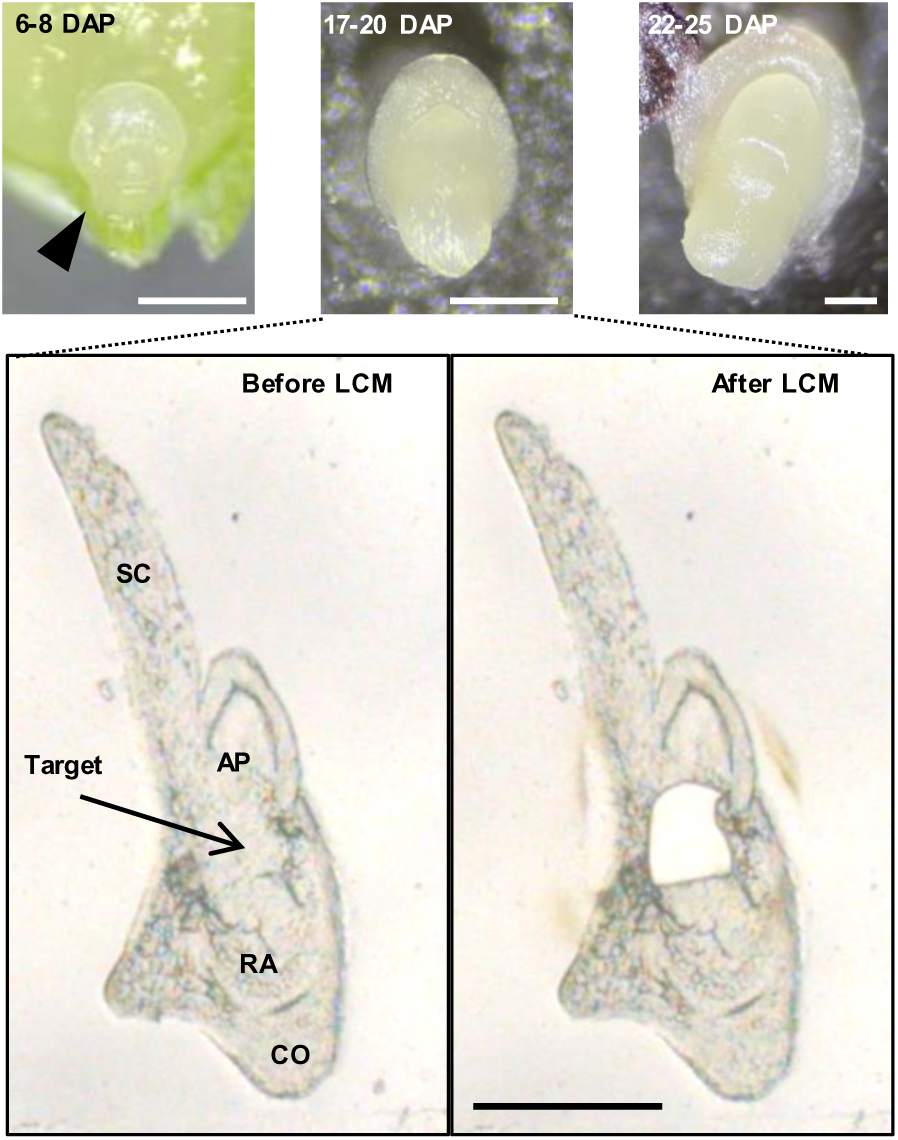
Embryo developmental stages and laser capture microdissection of the embryonic root for RNA extraction. Developmental stages of *Ae. speltoides* embryos used for RNA extraction. Top panels: representative embryos at 6–8 days after pollination (DAP), 17–20 DAP, and 22–25 DAP. Bottom panels: tissue cryosection (16 μm-thick) of a mid-stage embryo (17 DAP) before (left) and after (right) laser capture microdissection (LCM). The embryonic root targeted for RNA isolation is indicated by the arrow (Target). Anatomical landmarks: SC, scutellum; AP, apical meristem; RA, radicle; CO, coleorhiza. Scale bar: 400 μm.

**Supplementary Fig. 6.**
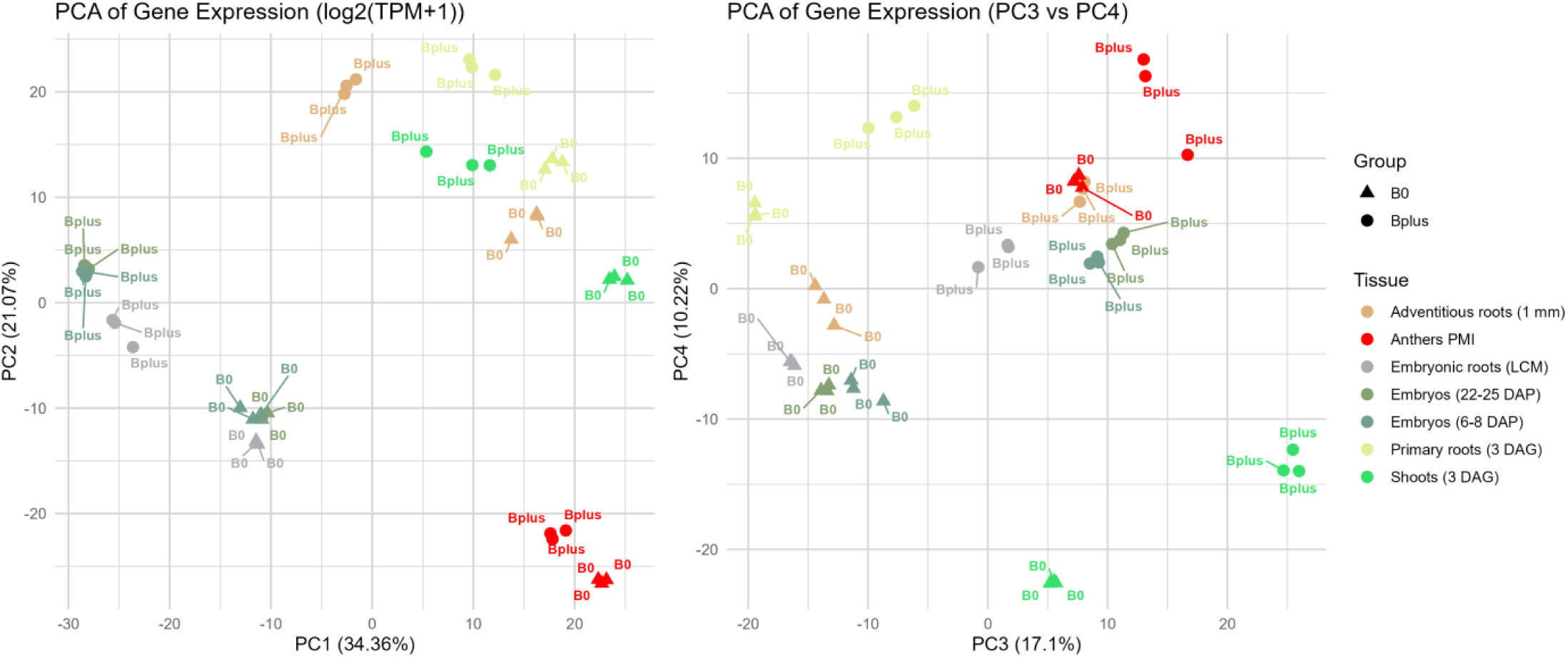
Global gene expression variation is primarily driven by tissue identity. PCA of gene expression profiles (log2(TPM+1)) across all 42 RNA-seq libraries. Left: PC1 vs PC2; Right: PC3 vs PC4. Shape indicates B chromosome status (triangle: 0B; circle: +B); color indicates tissue type.

**Supplementary Fig. 7.**
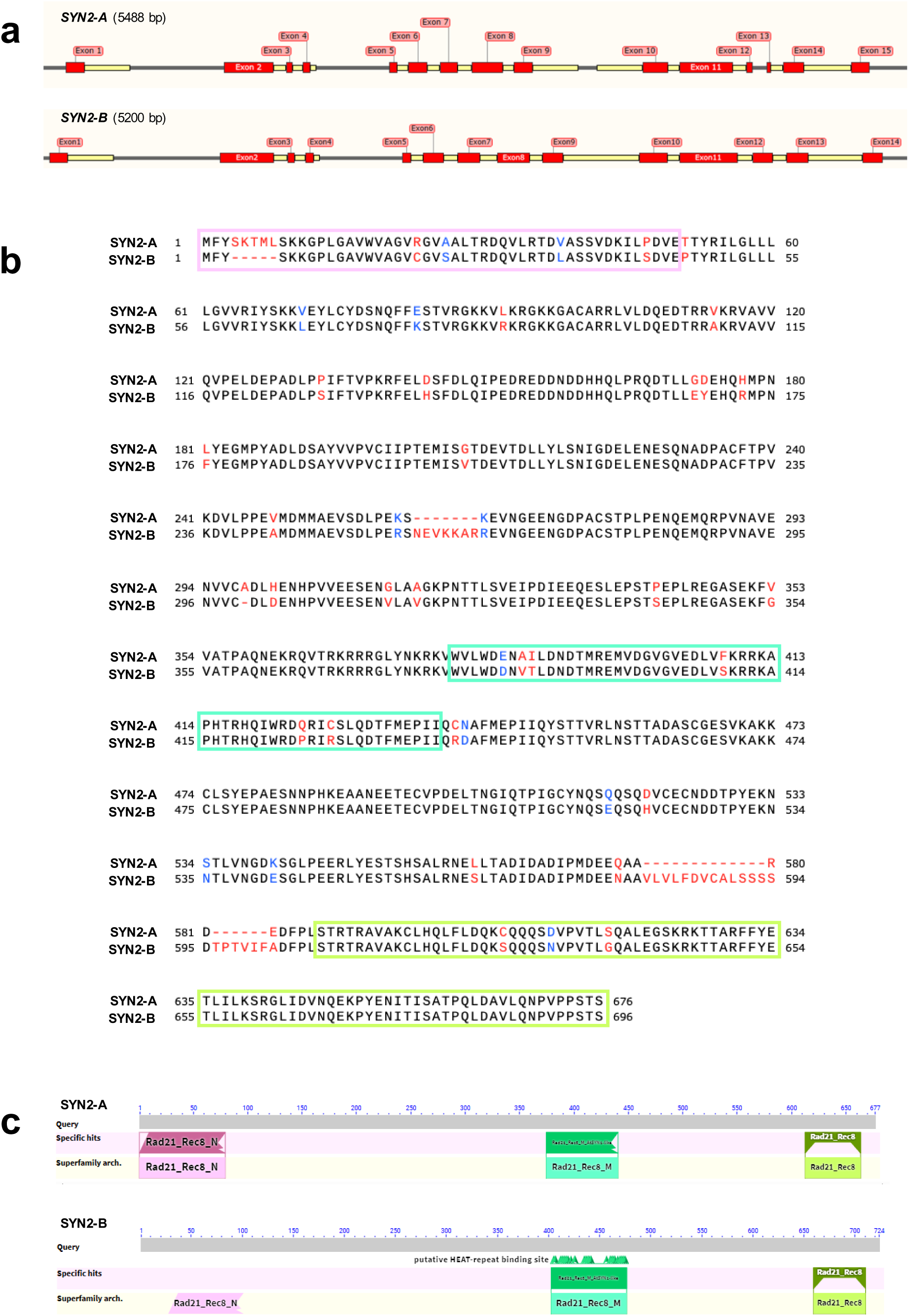
Structural comparison of SYN2-A and SYN2-B. (**a**) Gene structure of *SYN2-A* (5488 bp) and *SYN2-B* (5200 bp). Red boxes indicate exons. (**b**) Protein sequence alignment of SYN2-A and SYN2-B. Identical residues are shown in black; differences are highlighted in red, and conservatively substituted residues in blue. Numbers indicate amino acid positions. Colored boxes demarcate the three higher-confidence structural motifs of SYN2: the N-terminal RAD21/REC8 domain (pink), the central HEAT-repeat-binding (middle) motif (green), and the C-terminal RAD21/REC8 domain (yellow-green), corresponding to the regions confidently predicted by AlphaFold3 (Fig. 4c). (**c**) Domain architecture of SYN2-A and SYN2-B based on superfamily annotation. The conserved RAD21/REC8 cohesin domain and HEAT repeat-binding domain are indicated.

**Supplementary Fig. 8.**
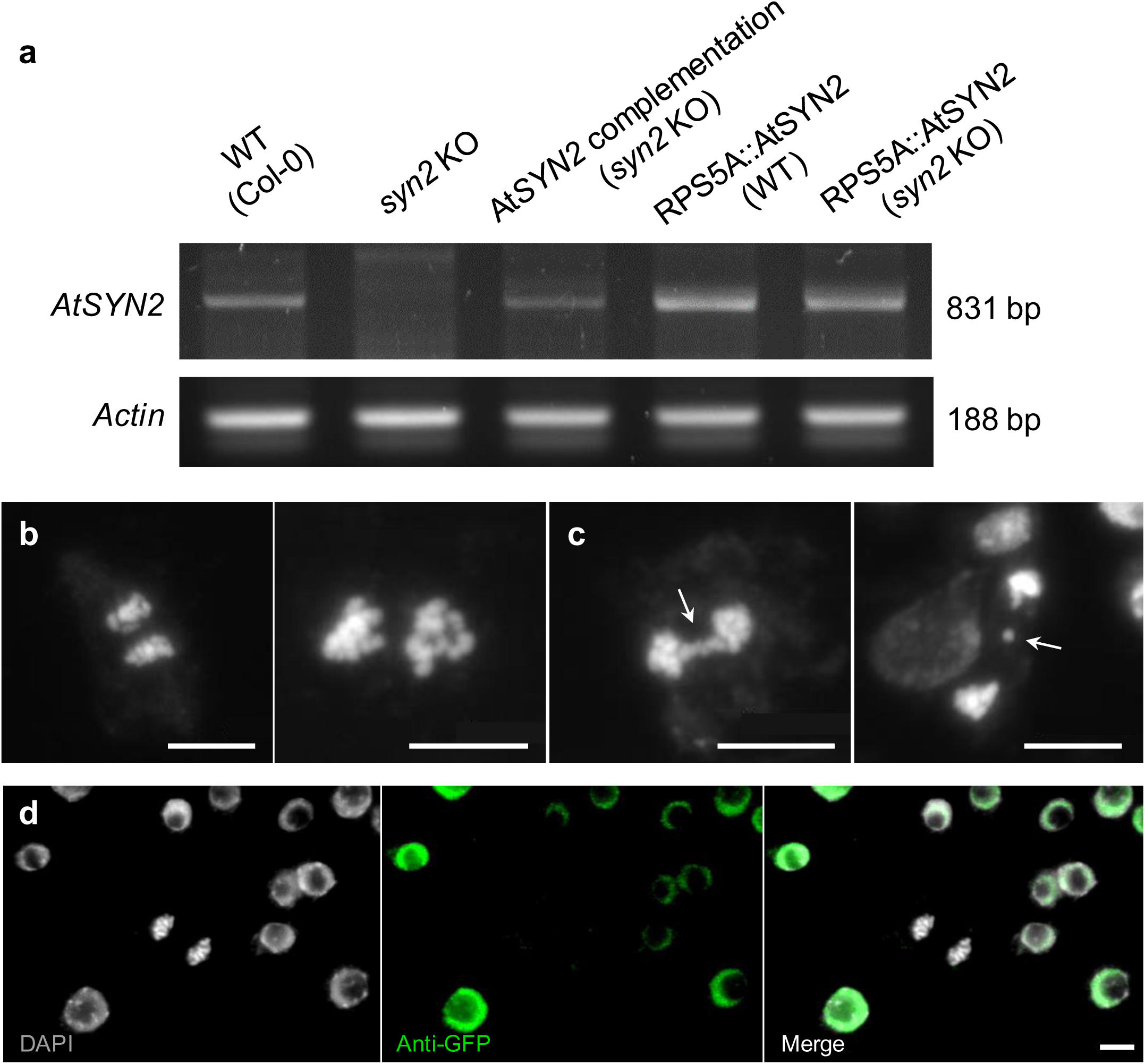
AtSYN2 expression validation and chromosome segregation phenotypes in *A. thaliana* loss-of-function and overexpression lines. (**a**) RT-PCR validation of *AtSYN2* transcript accumulation in wild-type Col-0, homozygous *syn2* knockout (T-DNA SALK_044851), *AtSYN2* complementation line in the *syn2* background (pAtSYN2::gAtSYN2-GFP), and RPS5A-driven overexpression lines in wild-type and *syn2* backgrounds (pRPS5A::GFP-AtSYN2). Expected amplicon sizes: *AtSYN2*, 831 bp; *actin*, 188 bp. Actin served as an internal control for RNA integrity and cDNA synthesis. (**b**) Representative DAPI-stained late anaphase in wild-type Col-0 showing precise chromosome segregation. (**c**) Representative DAPI-stained late anaphase in the pRPS5A::GFP-AtSYN2 line (wild-type background) showing chromosome segregation defects including a chromosome bridge and a lagging chromosome (arrows). (**d**) Anti-GFP immunostaining of pRPS5A::GFP-AtSYN2 root tips confirming GFP-AtSYN2 expression in interphase nuclei. Scale bars: 5 μm.

**Supplementary Fig. 9.**
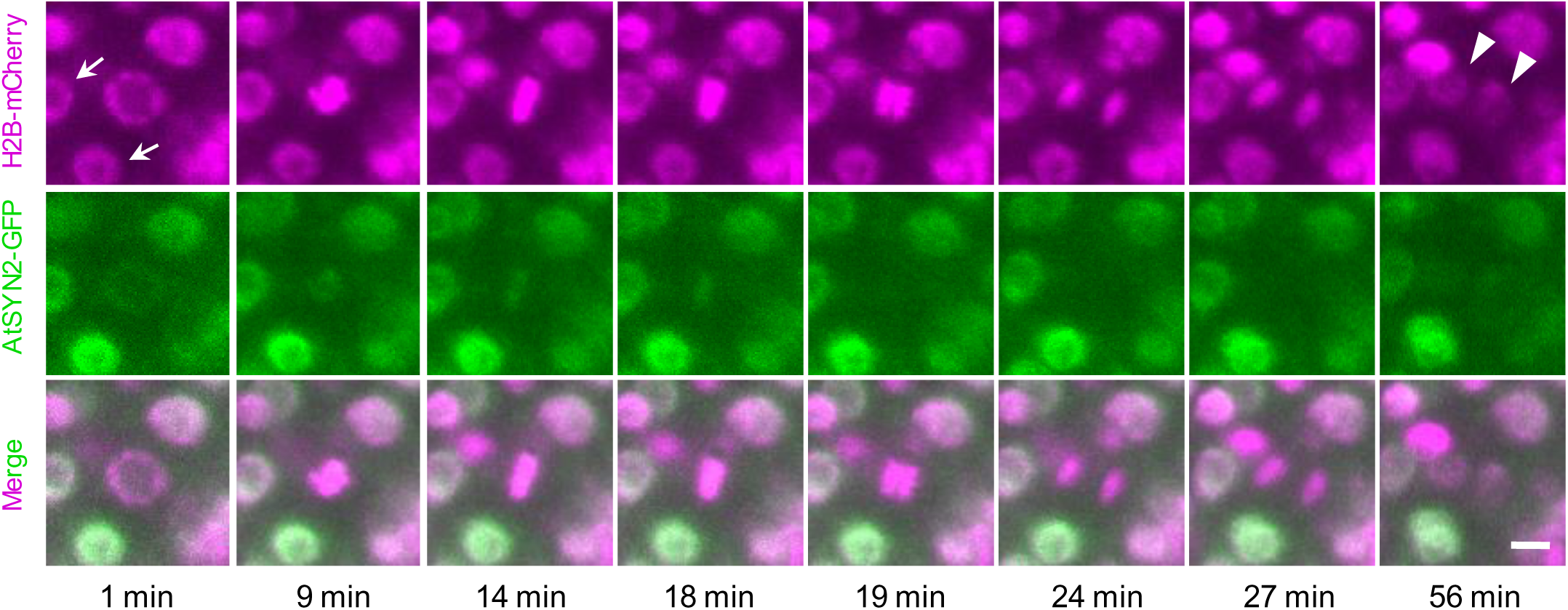
Mitotic dynamics of AtSYN2 under the control of the endogenous ATSYN2 promoter. Time-lapse images of a dividing *A. thaliana* root tip cell expressing pAtSYN2::AtSYN2-GFP (green) together with the chromatin marker H2B-mCherry (magenta). Arrows indicate interphase nuclei in surrounding cells at the onset of imaging. AtSYN2-GFP signal co-localizes with chromatin from interphase through metaphase and disappears at anaphase onset, consistent with normal cohesin release dynamics. Arrowheads indicate two daughter nuclei at the end of cell division. Scale bar: 5 μm.

**Supplementary Fig. 10.**
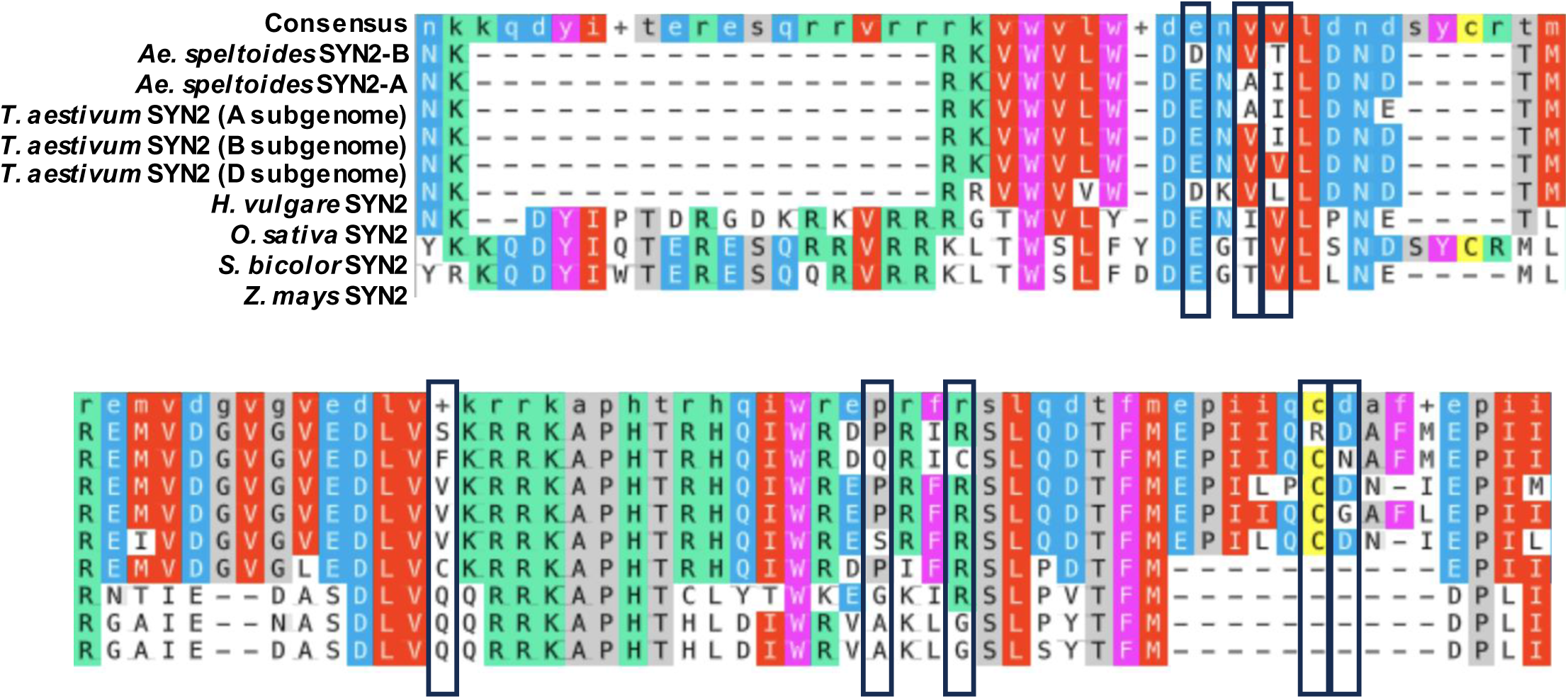
Cross-species sequence conservation of the SCC3-interacting region of SYN2. Multiple sequence alignment of the SCC3-binding region of SYN2 orthologs across cereals and other monocots, comprising the *Ae. speltoides* B- and A-encoded copies (SYN2-B and SYN2-A), the A, B, and D subgenome copies of *Triticum aestivum*, *Hordeum vulgare* subsp. *vulgare*, *Oryza sativa*, *Sorghum bicolor*, and *Zea mays*. The consensus sequence is shown on top, and residues are colored by physicochemical property. Black boxes indicate positions at which SYN2-B carries an amino acid substitution relative to SYN2-A; at these positions the residue is otherwise conserved across the SYN2 orthologs, indicating that several SYN2-B-specific substitutions occur at evolutionarily conserved sites within the SCC3-interacting interface.

**Supplementary Fig. 11.**
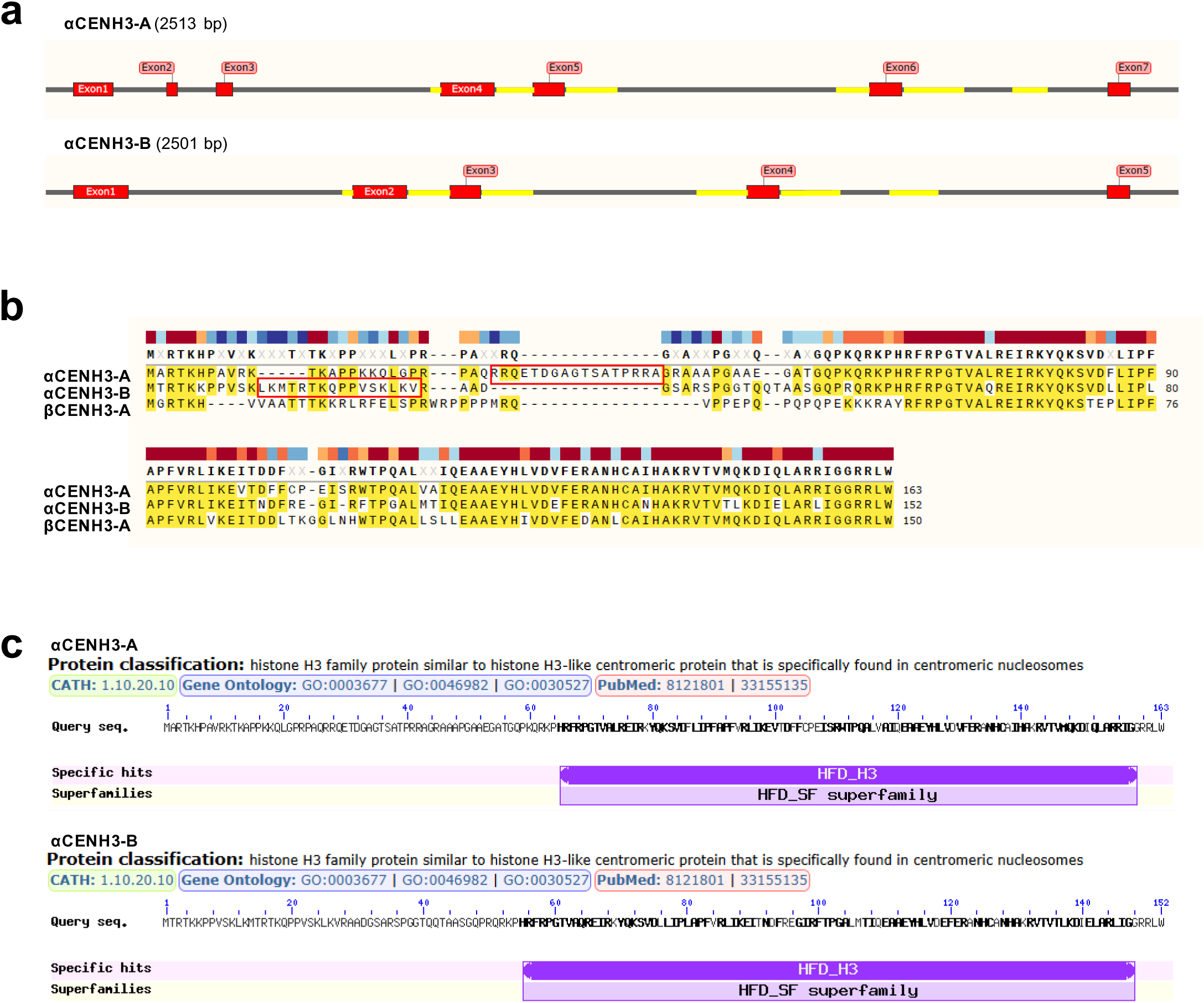
Structural and sequence comparison of αCENH3-A and αCENH3-B of *Ae. speltoides*. (**a**) Gene structure of *αCENH3-A* (2,513 bp) and *αCENH3-B* (2,501 bp), showing exon–intron organization. Conserved intron regions are highlighted in yellow. (**b**) Multiple sequence alignment of αCENH3-A, βCENH3-A, and αCENH3-B protein sequences. Conserved residues are highlighted in yellow. The N-terminal peptide sequences used for antibody production are indicated by red boxes. The histone fold domain is highly conserved across all three variants, whereas the greatest divergence is observed in the N-terminal tail. (**c**) Protein classification and domain assignment of αCENH3-A and αCENH3-B based on the NCBI conserved domain database. Both proteins are classified within the HFD_H3 superfamily, consistent with their identity as centromere-specific histone H3 variants.

**Supplementary Fig. 12.**
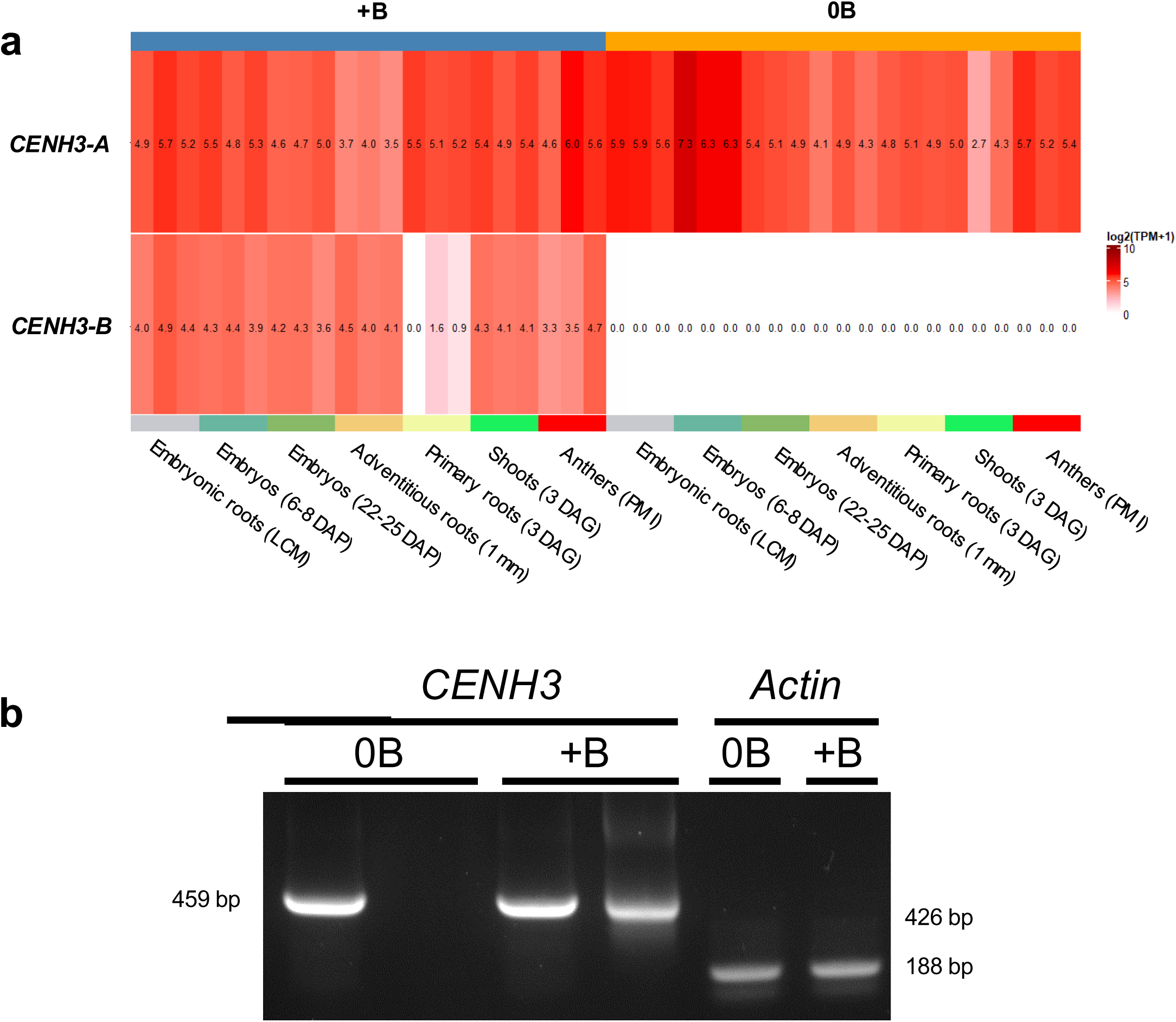
Expression profiles of CENH3-A and CENH3-B across *Ae. speltoides* tissues. (**a**) Heatmap of *CENH3-A* and *CENH3-B* expression (log2(TPM+1)) across 42 RNA-seq samples from 0B and +B plants spanning seven tissue types. *CENH3-A* is constitutively expressed across all tissues regardless of B’s presence. *CENH3-B* expression is restricted to +B tissues, with notably elevated levels in elimination-active tissues, shoots, and PM I anthers. (**b**) RT-PCR validation of *CENH3-A* and *CENH3-B* expression in 0B and +B PM I anther samples using gene-specific primers. Expected amplicon sizes: *CENH3-A*, 459 bp; *CENH3-B*, 426 bp. *Actin* (188 bp) served as an internal control for RNA integrity.

**Supplementary Fig. 13.**
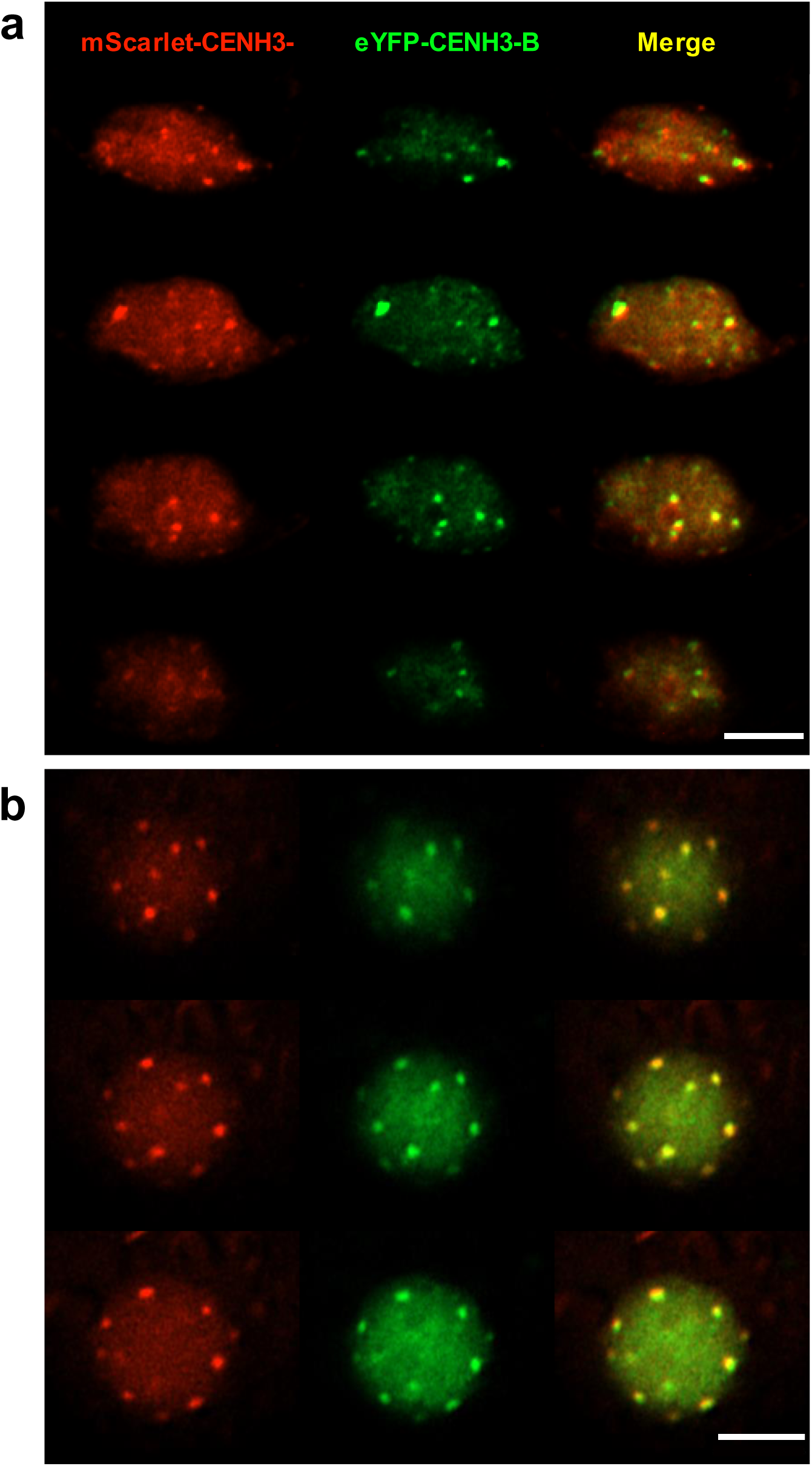
| **Co-localization of CENH3-A and CENH3-B signals in *Ae. speltoides* leaf protoplast nuclei.** Z-stack confocal series of a single nucleus from (**a**) a 0B protoplast and (**b**) a +B protoplast co-transformed with mScarlet-CENH3-A (red) and eYFP-CENH3-B (green). Both variants localize to the nucleus and form distinct punctate signals (14 signals for CENH3-A, 17 for CENH3-B). Merged images show colocalization of CENH3-A and CENH3-B signals irrespective of B chromosome presence, indicating that CENH3-B centromeric incorporation does not require B chromosome DNA. Scale bars: 5 μm.

**Supplementary Movie 1 |** Light sheet fluorescence microscopy was used to follow mitosis in a root meristem cell of an *Arabidopsis thaliana* pRPS5A::GFP-AtSYN2 line. GFP-AtSYN2 (green) marks the cohesin subunit and H2B-mCherry (magenta) marks chromatin. A chromosome bridge forms at anaphase onset, and the lagging chromatin is retained as a micronucleus at the end of mitosis. Selected frames are shown in Fig. 3d.

**Supplementary Movie 2 |** Light sheet fluorescence microscopy was used to follow mitosis in a root meristem cell of an *A. thaliana* pAtSYN2::AtSYN2-GFP line. AtSYN2-GFP (green) marks the cohesin subunit and H2B-mCherry (magenta) marks chromatin. The AtSYN2-GFP signal disappears at anaphase onset and sister chromatids separate without bridges or lagging chromatin. Selected frames are shown in Supplementary Fig. 9.

